# Response of total (DNA) and metabolically active (RNA) microbial communities in *Miscanthus × giganteus* cultivated soil to different nitrogen fertilization rates

**DOI:** 10.1101/2021.10.28.466385

**Authors:** Jihoon Yang, Jaejin Lee, Jinlyung Choi, Lanying Ma, Emily Heaton, Adina Howe

## Abstract

*Miscanthus* x *giganteus* is a promising high-yielding perennial plant to meet growing bioenergy demands but the degree to which the soil microbiome affects its nitrogen cycling and subsequently, biomass yield remains unclear. In this study, we hypothesize that contributions of metabolically active soil microbial membership may be underestimated with DNA-based approaches. We assessed the response of the soil microbiome to nitrogen availability in terms of both DNA and RNA soil microbial communities from the Long-term Assessment of Miscanthus Productivity and Sustainability (LAMPS) field trial. DNA and RNA were extracted from 271 samples, and 16S SSU rRNA amplicon sequencing was performed to characterize microbial community structure. Significant differences were observed in the resulting soil microbiomes and were best explained by the sequencing library of origin, either DNA and RNA. Similar numbers of taxa were detected in DNA and RNA microbial communities, with more than 90% of taxa shared. However, the profile of dominant taxa within DNA and RNA differed, with varying proportions of *Actinobacteria* and *Proteobacteria* and *Firmicutes* and *Proteobacteria*. Only RNA microbial communities showed seasonal responses to nitrogen fertilization, and these differences were associated with nitrogen-cycling bacteria. The relative abundance of bacteria associated with nitrogen cycling was 7-folds higher in RNA than in DNA, and genes associated with denitrifying bacteria were significantly enriched in RNA, suggesting that these bacteria may be underestimated with DNA-only approaches. Our findings indicate that RNA-based SSU characterization can be a significant and complementing resource for understanding the role of soil microbiomes in bioenergy crop production.

**Importance:** *Miscanthus* x *giganteus* is becoming a cornerstone of bioeconomy cropping systems, but it remains unclear how the soil microbiome supplies nitrogen to this low-input crop. DNA-based techniques are used to provide community characterization but may miss important metabolically active taxa. By analyzing both DNA- and actively transcribed RNA-based microbial communities, we found that nitrogen cycling taxa in the soil microbiome may be underestimated using only DNA-based approaches. Accurately understanding the role of microbes and how they cycle nutrients is important for the development of sustainable bioenergy crops, and RNA-based approaches are recommended as a complement to DNA approaches to better understand the microbial, plant, and management interactions.

## Introduction

The sterile allopolyploid (2n=3x=57) *Miscanthus × giganteus* (Greef et Deu.) is a promising perennial grass bioenergy crop because of its ability to produce large amounts of biomass with little fertilizer compared to hay or grain crops (1–4). The peak biomass production of *M.* x *giganteus* has been observed to be up to three times higher than switchgrass (*Panicum virgatum* L. cv. Cave-in-Rock), similar to willow (*Salix schwerinii* E. Wolf × viminalis L.), three times higher than reed canary grass (*Phalaris arundinacea* L.), and two times higher than triticale (*Triticosecale* Wittmack) (5–7). Additionally, *M.* x *giganteus* production has been shown to have decreased environmental impact, with decreased requirements of nitrogen and pesticides (8, 9) and reduced nitrate leaching relative to other bioenergy crops (10, 11). These advantages of *M.* x *giganteus* and its ability to maintain high productivity for up to 20 years compared to other energy crops have contributed to its increased cultivation (9, 12–14).

To support its growth, environmental and management factors that can affect the productivity of *M.* x *giganteus* have been evaluated. Previously, *M.* x *giganteus* has been observed to decrease in productivity at low temperatures (15, 16). It has also been observed to have relatively high water demand (17, 18) and to require cultivation for at least three years to obtain adequate yield (16, 19–25). Recommendations for nitrogen fertilization of *M.* x *giganteus* are inconsistent, with previous studies showing that fertilization can have little to no effect (26–31) or contribute to its productivity (32–35).

Previously, it has been estimated that *M.* x *giganteus* can obtain 16% of its nitrogen demand from the atmosphere during the growing season (36). Nitrogen can also be provided by the activity of nitrogen-fixing bacteria in the rhizobiome of *M.* x *giganteus* (37), which are enriched early after *M.* x *giganteus* planting (36). Nitrogen fixation genes have been observed to be more abundant in *M.* x *giganteus* relative to other energy crops planted in similar soils (38, 39). Specific phyla which have been identified in *M.* x *giganteus* rhizobiomes include *Actinobacteria* and *Proteobacteria*, which include known nitrogen-fixing families such as *Hyphomicrobiaceae*, *Bradyrhizobiaceae*, *Rhodospirillaceae*, and *Geobacteraceae* (40).

To date, all studies of *M.* x *giganteus* soil microbial communities and their response to fertilization or biomass production have been limited to the characterization of soil environmental DNA. We previously used sequencing of 16S rRNA genes in DNA to identify significant interactions between microbial diversity, stand age, fertilization, and above-ground biomass in *M.* x *giganteus* (41). However, it is possible that DNA-based analysis may underestimate the number of active taxa, resulting in biased interpretations of how microbial communities respond to the environment (42, 43). By contrast, RNA-based characterization of microbial communities, representing metabolically active or transcribed genes, can better relate community responses to environmental variability (44–48). Additionally, RNA-based studies are more sensitive and have detected underrepresented active bacteria that are below the amplification threshold of DNA-based approaches. Despite the advantages of RNA-based methods, direct comparison of the DNA and RNA methods for microbial community characterization in bioenergy crops soil microbial communities is sparse. One previous study of the bioenergy grass, *Pennisetum purpure*, compared bacterial communities of DNA- and RNA-based denaturing gradient gel electrophoresis (DGGE) profiles and clone libraries and found that RNA-based methods could identify enriched metabolically active membership (49).

In this study, we perform comparison of DNA and RNA approaches to help us better understand how soil microbiome in field-grown *M.* x *giganteus* can inform management and environmental impacts of *M.* x *giganteus* production. We evaluate the effects of stand age (representing different initial growth environments) and fertilization (representing different N availability) on changes in microbial community membership and structure. We hypothesize that microbiome responses (as indicated by DNA and RNA) to *M.* x *giganteus* management will differ and specifically that metabolically active (RNA) microbial communities will show a more rapid and sensitive response to fertilization than total (DNA) microbial communities. To test these hypotheses, soil samples were collected from the Long-term Assessment of Miscanthus Productivity and Sustainability (LAMPS) site, a replicated chronosequence field previously used to investigate the effects of stand age and nitrogen fertilizer on *M.* x *giganteus* and corn (*Zea mays* L.) (28, 50). DNA and RNA were extracted from these soil samples, and we compared these microbial responses to stand age, N fertilization amount, and time since fertilization.

## Material and Methods

### Sample description

Soil samples were collected in 2018 from the Long-term Assessment of Miscanthus Productivity and Sustainability (LAMPS) site located in Central IA, USA (42.013° N, 93.743° W). This staggered-start experiment was planted with *M.* x *giganteus* (clone “Freedom”, AGgrow Tech, High Point NC, USA) at a density of ∼11 plants m^-2^ in replicated blocks (n=4) in each of 2015, 2016 and 2017 as described previously (28). The experimental design is a split-plot replicated block with age (planting year) as the main plot and N fertilization rate as the split plot. Soils at the site are deep loams (>1m) formed over glacial till; the dominant soil type (53%) is a Webster clay loam (fine-loamy, mixed, superactive, mesic Typic Endoaquoll). Fertilizer was applied as banded urea ammonium nitrate (UAN) in aqueous solution and side-dressed into the soil at 0.1 m depth on May 9, 2018, at rates of 0, 224, and 448 kg ha^-1^ N. Soil samples were taken on April 30, May 14, May 30, and July 3. Soils were collected from within 10 cm radius of the *M.* x *giganteus* stems using a sampling core (30.5 cm wet sample tube with 1.75cm diameter, Clements Associates Inc, USA). Soil samples included in this analysis were obtained in triplicate from 60 experimental plots at each time point and analyzed independently. Samples for DNA extraction were stored on dry ice immediately after being taken as described previously (41), and samples for RNA extraction were immediately collected and then frozen in RNAlater (Thermo Fisher Scientific, USA) which offers the advantage of preserving microbial community integrity while preventing RNA degradation (51). All samples were stored in a cooler filled with dry ice during return to the laboratory.

### DNA/RNA extraction and 16S rRNA gene amplicon sequencing

DNA and RNA extraction was performed from subsampled 0.25 g soil samples submerged in RNAlater (Thermo Fisher Scientific, USA), using the MagAttract PowerMicrobiome DNA/RNA EP Kit (Qiagen, USA) following the standard protocol in this kit and liquid handling in Eppendorf epMotion 5075 (Eppendorf North America). The extracted RNA was transcribed into cDNA according to a standard protocol using iScript™ cDNA Synthesis Kit (BIO-RAD, USA) for sequencing analysis. The resulting DNA and RNA were analyzed for quantity using an Invitrogen™ Qubit™ 4 Fluorometer (Invitrogen, USA). DNA and RNA sample concentrations above 10 ng ul^-1^ were normalized to 10 ng ul^-1^ prior to sequencing. Samples with concentrations lower than 10 ng ul^-1^ were submitted directly for sequencing. The V4 region of the bacterial 16S rRNA gene was amplified with the conserved primers 515F (5′-GTGYCAGCMGCCGCGGTAA-3′) and 806R (5′-GGACTACNVGGGTWTCTAAT-3′) (52, 53). Bacterial amplicon sequencing was performed on Illumina Miseq with Miseq Reagent Kit V2 (Illumina, USA) at Argonne National Laboratory. The DNA and RNA sequencing data are available at NCBI Short Read Archive PRJNA601860 and PRJNA745191, respectively.

### Amplicon bioinformatics and statistical analysis

The *DADA2* package (version 1.13.1) in R (version 4.1.0) was used to perform quality control of sequencing libraries and to determine the abundance of amplicon sequence variants (ASV). The quality filtering parameters for all sequences were the same as previously described for DNA amplicons (41). The Ribosomal Database Project (RDP) Classifier (version 11.5) was used for taxonomic identification of each observed ASV depending on the sequence similarity to the representatives in the current database. ASVs were removed if no more than 10 total observations were observed in a sample. All statistical analyses were performed in R (version 4.1.0). Two diversity indices, Chao1 and Shannon, were used to compare the alpha diversity of bacteria using the *vegan* package (version 2.5 - 7). Multivariate homogeneity of group dispersions, calculating the average distance of members to the centroid of the group, was used to analyze the dispersion of each sample using *betadisper* function from the *vegan* package (version 2.5 - 7). Significant differences in alpha diversity and homogeneity between DNA and RNA microbial communities were evaluated using the Kruskal-Wallis test with Dunn’s post hoc test. Permutational multivariate analysis of variance (PERMANOVA) was performed with the *adonis* function of the *vegan* package using the Bray-Curtis dissimilarity matrix (version 2.5 - 7). PERMANOVA was performed to identify significant differences between centroids of each microbial community, and the R^2^ statistic represents the proportion of the variance for the separation of the microbial community that was explained by experimental and field environmental factors (i.e., origin of the sequencing library, stand age, N fertilization amount, fertilization history, and time since fertilization). PERMANOVA was performed using the “strata” argument for the planted block, which was identified as one of the major factors to structure the microbial composition in the previous study, to better identify the effects of stand age and N fertilization amount, fertilization history, and time since fertilization. This analysis restricted permutations to the dataset within each block and was used to quantify variations between and within treatments (41). The comparison between the two groups within the three (stand age, N fertilization amount) or four (time since fertilization) groups was accomplished using pairwise PERMANOVA. The level of significance in the statistical analysis was defined as p < 0.05.

## Results

### DNA and RNA microbial communities differ in microbial composition and alpha diversity

The 16S rRNA amplicons from DNA and RNA from soil samples representing three ages and three fertilization rates of *M* x *giganteus* were compared. The origin of the sequencing library, either DNA or RNA, was found to have the greatest influence on the separation of the microbial community (R^2^_PERMANOVA_ = 0.117, p_PERMANOVA_ = 0.001, Table S1). Their influence was 3.9 times higher than that of stand age (R^2^_PERMANOVA_ = 0.030, p_PERMANOVA_ = 0.001) and 15 times higher than that of N fertilization amount (R^2^_PERMANOVA_ = 0.008, p_PERMANOVA_ = 0.001). DNA and RNA microbial communities were observed to separate into clear clusters using constrained analysis of principal coordinates (CAP) along the first axis of the CAP (CAP1, 11.8%, F = 77.56, p_ANOVA_ = 0.001, Figure 1). The microbial composition (ADONIS, p_ADONIS_ = 0.001) and homogeneity (betadisper, p_betadisper_ = 0.001) of DNA and RNA communities were also observed to be significantly different.

**Figure 1.**
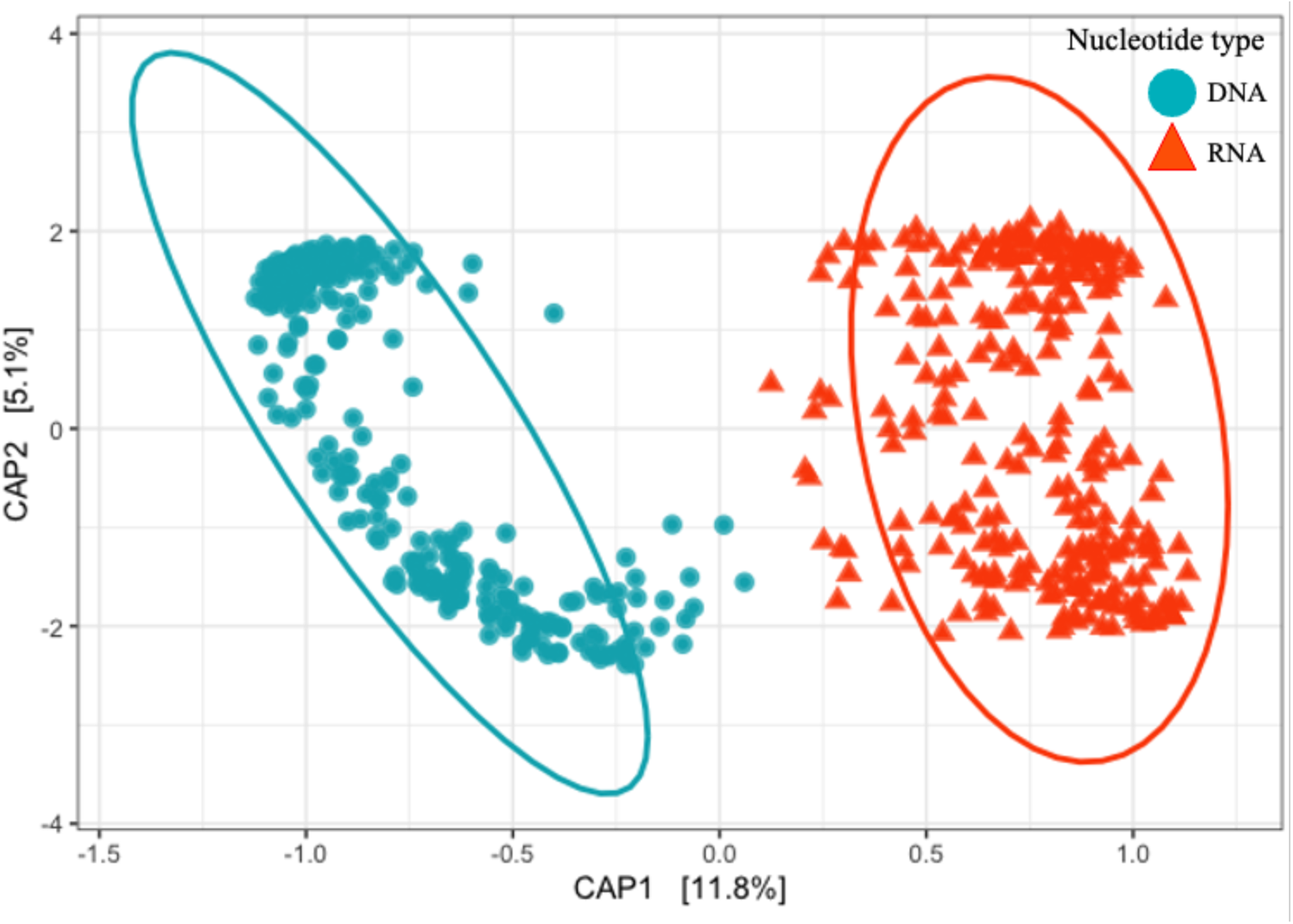
Similarities, assessed with Bray-Curtis indices, between DNA and RNA microbial communities from *M.* x *giganteus* soils. Constrained analysis of principal coordinates (CAP) was used to ordinate Bray-Curtis indices calculated with ASVs. Blue dot and red triangle represent the DNA and RNA microbial communities, respectively.

Alpha diversity of soil microbial communities was compared using the Shannon index, which evaluates both microbial richness and evenness, and Chao1, which evaluates the abundance of observed species. Both alpha diversity indices showed significant differences between DNA and RNA microbial communities, with higher alpha diversity observed in DNA microbial communities (p_Shannon_ < 0.001, p_Chao1_ < 0.001, Figure 2). On average 32% and 12% higher Chao1 and Shannon indices, respectively, were observed in DNA compared to RNA microbial communities. The average value of alpha diversity was higher in DNA, but the variation in alpha diversity indices was larger between RNA samples. Specifically, the DNA Chao1 index was in the range of 1,667 to 9,170, and RNA was associated with a much wider range of 195 to 9,343. Similar results were observed with Shannon indices, with DNA ranging from 4.88 to 7.85 and RNA from 2.69 to 7.72.

**Figure 2.**
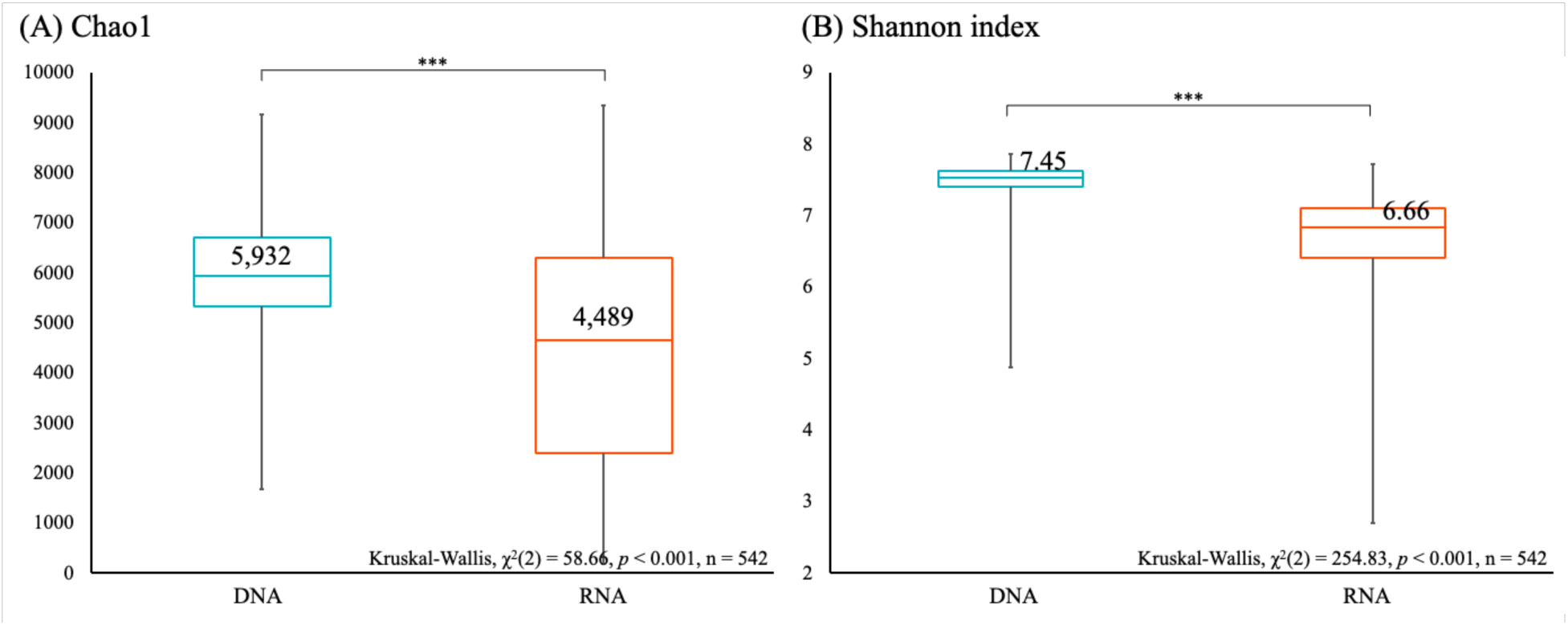
Alpha diversity indices of DNA and RNA microbial communities. Richness ((A) Chao1, (B) Shannon index) were estimated for microbial communities with ASVs. Letters “***” denote significant differences of alpha diversity indices between DNA and RNA microbial communities at a p-value < 0.05 as assess by Kruskal-Wallis with post hoc Dunn’s test.

### Taxa distributions varied between DNA and RNA microbial communities

The total number of taxa in DNA and RNA microbial communities was estimated by observations of ASVs, where a total of 39,898 and 32,171 ASVs were identified in DNA and RNA, respectively. We compared the ASVs between DNA and RNA samples and found that 17,779 ASVs were shared between DNA and RNA microbial communities (32% and 58%, respectively); 22,119 and 14,392 ASVs were unique in DNA and RNA, respectively. Unique ASVs were generally low abundance (average < 0.000003%) and low prevalence (average < 0.022%) in their respective libraries (Figure 3). ASVs that were identified in both DNA and RNA were found to be identified at increased though still low abundance (average < 0.00005%) and higher prevalence (average > 0.16%).

**Figure 3.**
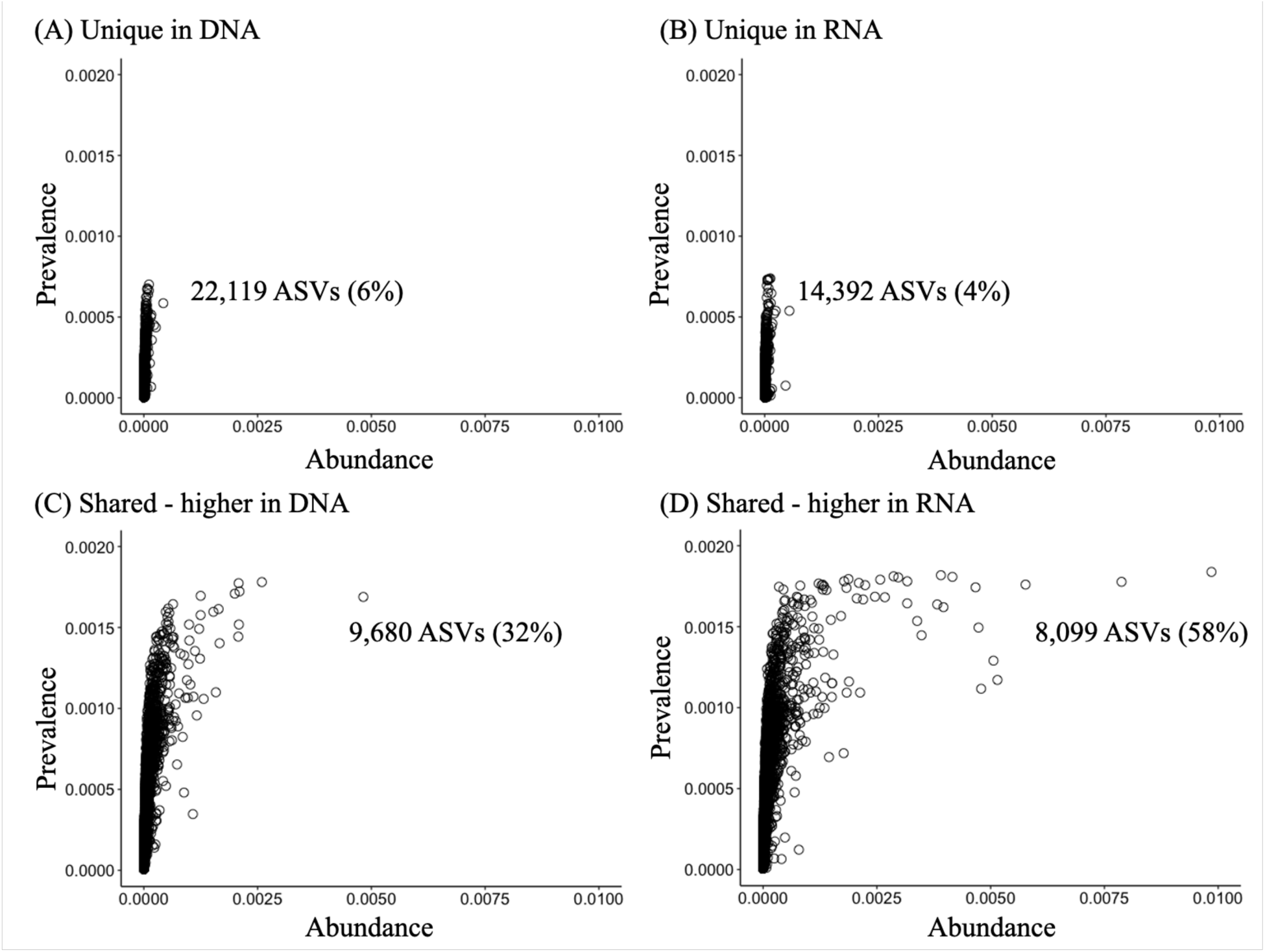
Abundance-occupancy comparison of ASVs in the DNA and RNA microbial communities. Abundance-occupancy distributions were assessed to identify the dynamics of the DNA and RNA microbial community memberships. Each point is an ASV. The ASVs were classified as (A) unique in DNA or (B) unique in RNA, respectively, when it was detected only in the DNA or RNA microbial communities. ASVs detected in both DNA and RNA microbial communities were classified as “shared” and further classified by the average relative abundance based on its enrichment in (C) DNA or (D) RNA.

ASVs commonly identified between DNA and RNA libraries were further classified based on their enrichment in DNA or RNA, specifically using the ratio of RNA:DNA relative abundances. The RNA:DNA ratio of shared ASVs ranged from 0.0023 to 1,300. The majority of shared ASVs (58%) were more enriched in RNA relative to DNA (Figure 4). For ASVs enriched in DNA (RNA:DNA ratio < 1), the average RNA:DNA ratio was 0.44; the average RNA:DNA ratio for ASVs enriched in RNA (RNA:DNA ratio > 1) was 4.82. Additionally, more variation was observed in shared ASVs which were enriched in RNA relative to those enriched in DNA.

**Figure 4.**
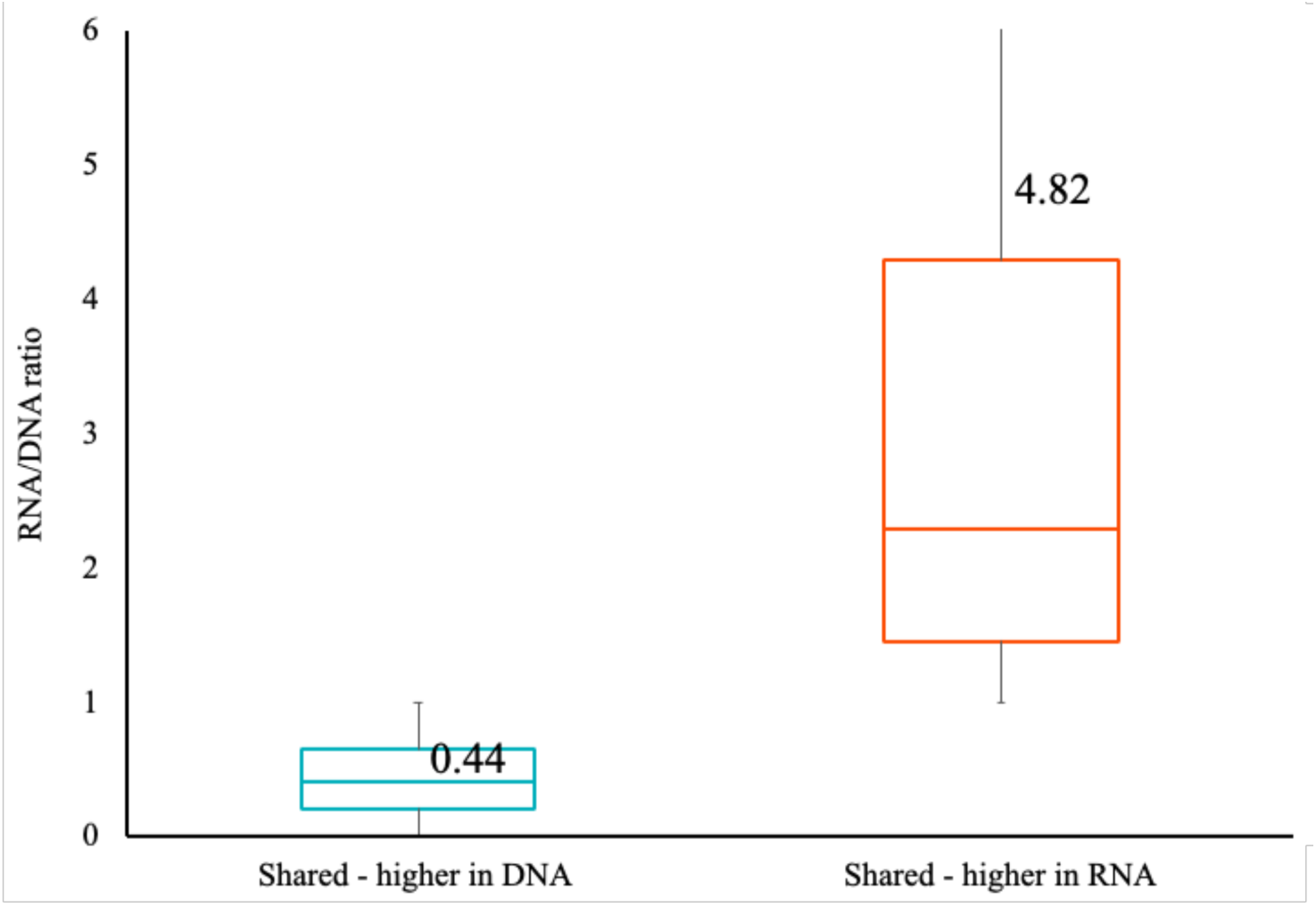
RNA/DNA ratio comparison of the shared ASVs. The ratio of average relative abundance in DNA and RNA microbial communities of ASVs detected in both microbial communities was compared to identify the biased in the DNA and RNA-based microbial community analysis results. Shared - higher in DNA (blue) and Shared - higher in RNA (red).

### Phylogenetic composition varied between DNA and RNA microbial communities

The phylogenetic composition of DNA and RNA microbial communities was compared, with 20 phyla identified in both libraries. Soils were dominated by *Actinobacteria* (26%) and *Proteobacteria* (33%) in DNA and mainly *Proteobacteria* (49%) in RNA (Figure 5). While DNA and RNA had similar membership at the phylum-level, the relative abundance of every phylum significantly differed (Table S2). Thirteen out of 20 phyla were more enriched in DNA than RNA, and seven phyla were more enriched in the RNA microbial communities.

**Figure 5.**
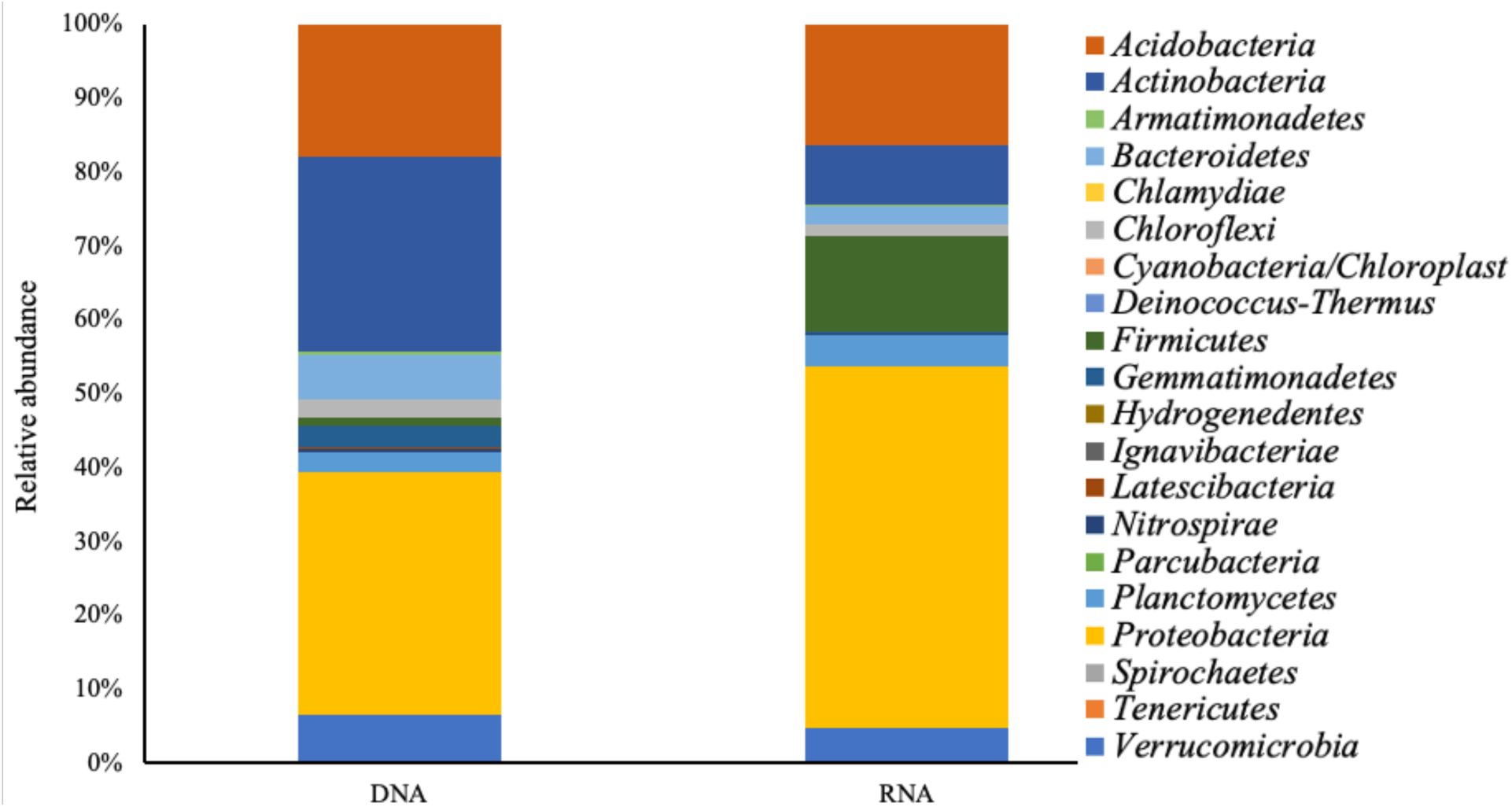
Phylum level differences in DNA and RNA microbial communities. Relative abundances of annotated ASVs are shown, identified to their closest match in the RDP classifier. See Supplementary Table 2 for a different perspective on the dynamics of numerical relative abundance.

We evaluated whether the phyla observed to be significantly different between DNA and RNA were comprised of ASVs unique to DNA or RNA or shared between the two methods (Figure S1). ASV shared by DNA and RNA microbial communities showed more pronounced variations in microbial community structures differences. *Actinobacteria* and *Bacteroidetes* were more enriched in DNA (p_Kruskal-Wallis_ < 0.05), while *Firmicutes* and *Proteobacteria* were more enriched in the RNA microbial community (p_Kruskal-Wallis_ < 0.05). Differentiating ASVs unique in DNA included sequences associated with *Actinobacteria*, *Gemmatimonadetes*, *Latescibacteria*, and *Parcubacteria*; in contrast, sequences associated with *Firmicutes* were unique in RNA.

### DNA and RNA microbial community compositions were variably changed by stand age, N fertilization amount, and time since fertilization

Previously, the response of the soil microbial community at this site to plant stand age, fertilization history, and time since fertilization was studied based on DNA (41). In this study, subsets of these samples were studied to directly compare DNA and RNA 16S rRNA gene characterization. Based on DNA, community composition responded significantly to the stand age and N fertilization amount. The community response based on RNA was similar, with the notable exception that time since fertilization showed a significant effect only in RNA (Table 1). The effect of stand age and N fertilization amount was generally larger in DNA than RNA, and time since fertilization had a larger effect on the RNA microbial community.

**Table 1.**
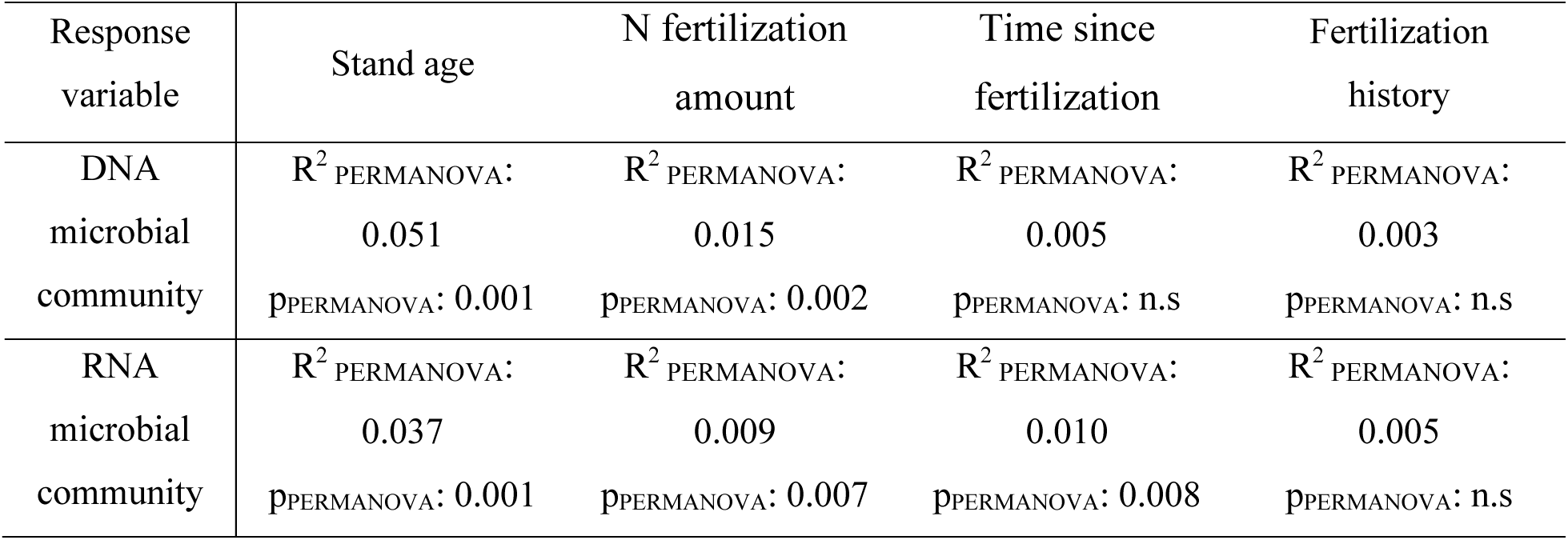
Permutational multivariate analysis of variance (PERMANOVA) for comparing DNA and RNA microbial community dissimilarity.

| Response variable | Stand age | N fertilization amount | Time since fertilization | Fertilization history |
| --- | --- | --- | --- | --- |
| DNA microbial community | $R^2_{\text{PERMANOVA}}$: 0.051<br>$p_{\text{PERMANOVA}}$: 0.001 | $R^2_{\text{PERMANOVA}}$: 0.015<br>$p_{\text{PERMANOVA}}$: 0.002 | $R^2_{\text{PERMANOVA}}$: 0.005<br>$p_{\text{PERMANOVA}}$: n.s | $R^2_{\text{PERMANOVA}}$: 0.003<br>$p_{\text{PERMANOVA}}$: n.s |
| RNA microbial community | $R^2_{\text{PERMANOVA}}$: 0.037<br>$p_{\text{PERMANOVA}}$: 0.001 | $R^2_{\text{PERMANOVA}}$: 0.009<br>$p_{\text{PERMANOVA}}$: 0.007 | $R^2_{\text{PERMANOVA}}$: 0.010<br>$p_{\text{PERMANOVA}}$: 0.008 | $R^2_{\text{PERMANOVA}}$: 0.005<br>$p_{\text{PERMANOVA}}$: n.s |

Next, pairwise comparisons of DNA and RNA microbial communities between stand ages were performed (pairwise PERMANOVA, Table S3). The stand ages of *M.* x *giganteus* included were 2-, 3-, and 4-year-old, and the microbial community of each stand age was significantly different based on both DNA and RNA microbial communities (p_pairwisePERMANOVA_ < 0.05). Similar patterns were observed for the response to N fertilization amount in both libraries, and both DNA and RNA microbial communities were found to have different microbial community compositions for three varying N fertilization amount (p_pairwisePERMANOVA_ < 0.05). Pairwise comparison of DNA and RNA based on sampling day or the time since fertilization resulted in no significant differences observed in DNA, but significant differences between pre-fertilization (10 days before fertilization) and 69 days since fertilization in RNA (Table S4, p_pairwisePERMANOVA_ < 0.05).

Stand age was consistently observed to explain the most variation between experimental factors, regardless of DNA or RNA methods (Table 1). We next evaluated if the specific phyla found to be different between stand ages was consistent between DNA and RNA microbial communities. A total of 20 identical phyla were detected in both sequencing libraries. Every phylum showed significant relative abundance differences between DNA and RNA (p_Kruskal-Wallis_ < 0.05). The dominant phyla differed between DNA and RNA (Figure 6), with *Acidobacteria* (>17%), *Actinobacteria* (>24%), and *Proteobacteria* (>32%) dominant in DNA, and *Firmicutes* (>12%) and *Proteobacteria* (>43%) in RNA. We subsequently selected these phyla to evaluate genera level differences between DNA and RNA methods.

**Figure 6.**
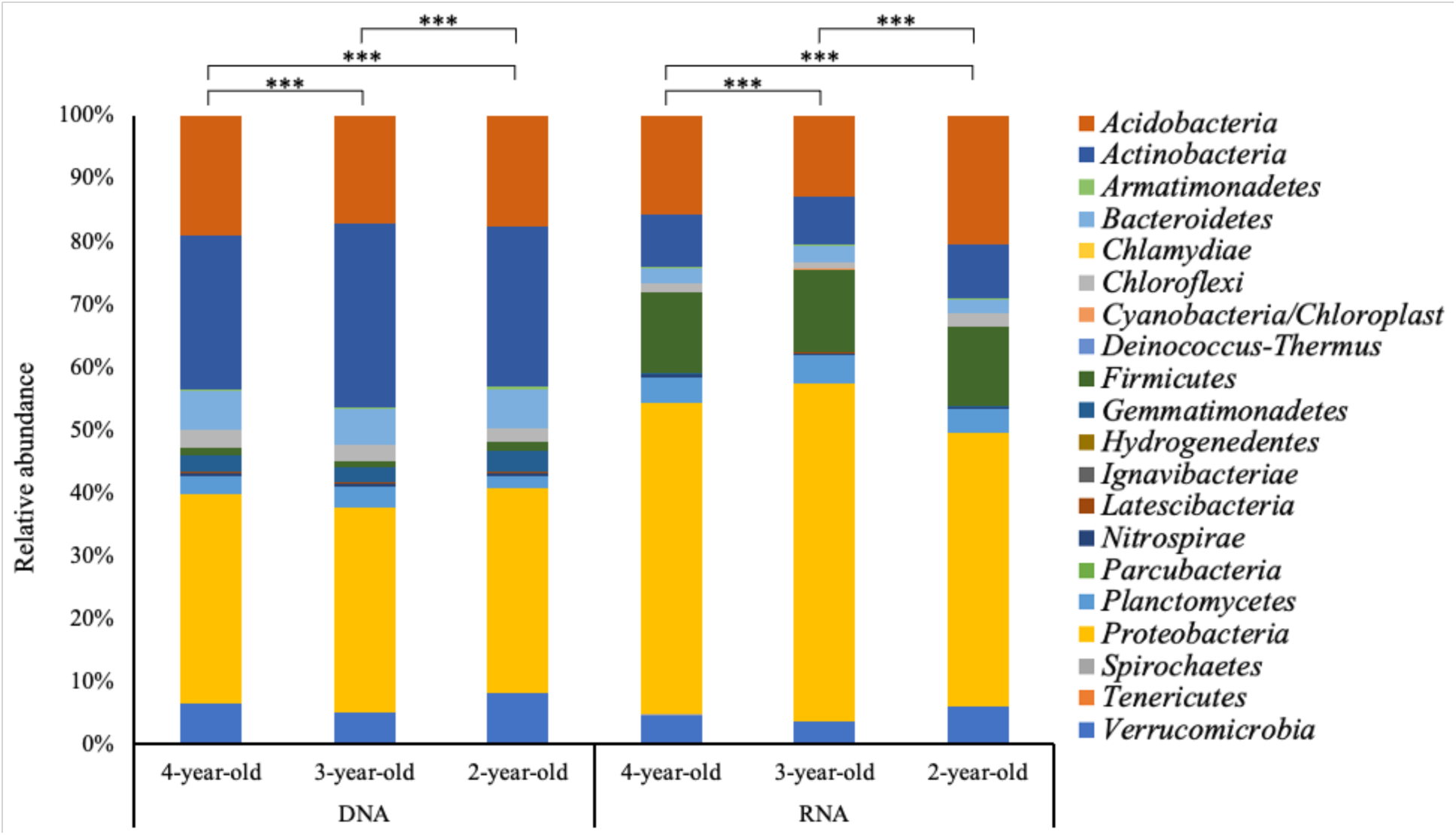
Phylum level differences in DNA and RNA microbial communities according to stand age differences. Relative abundances of annotated ASVs are shown, identified to their closest match in the RDP classifier. The letters “***” denote significant differences of relative abundance between different stand ages of *M.* x *giganteus* at a p-value < 0.05 as assessed by Kruskal-Wallis with post hoc Dunn’s test.

A total of 569 genera were detected among *Actinobacteria*, *Proteobacteria,* and *Firmicutes* and 308, 316, and 337 genera in 2-, 3-, and 4-year-old *M.* x *giganteus*, respectively, showed significant differences between the DNA and RNA microbial communities. We selected the genera with greater than 0.1% relative abundance and compared differences between taxonomic profiles in DNA and RNA (Figure S2). Sequences associated with *Bacillus*, *Clostridium*, *Paenibacillus*, *Sporosarcina* of *Firmicutes* and *Bradyrhizobium*, *Methyloversatilis*, *Nitrosomonas*, *Nitrosospira*, and *Steroidobacter* of *Proteobacteria* were more enriched in RNA than in DNA. On the other hand, *Gaiella* and *Solirubrobacter* of *Actinobacteria* were more enriched in DNA.

Differences in response to fertilization were also observed between DNA and RNA microbial communities. Both DNA- and RNA-based methods identified that soil microbial communities showed different responses to N fertilization amount (Table 1, Table S3), though the phylogenetic profile observed under fertilized conditions differed based on the two methods (Figure S3). Overall, a greater number of phyla in RNA relative to DNA were significantly affected by differences in N fertilization amount (Table S5). Seven phyla in RNA (*Actinobacteria*, *Firmicutes*, *Gemmatimonadetes*, *Hydrogenedentes*, *Latescibacteria*, *Nitrospirae*, and *Proteobacteria*) showed significant differences between N fertilization amount differences compared to four phyla in DNA (*Acidobacteria*, *Chloroflexi*, *Latescibacteria*, and *Proteobacteria*). *Actinobacteria* (>26%) was more enriched in DNA microbial communities (p_Kruskal-Wallis_ < 0.05), and *Firmicutes* (>12%) and *Proteobacteria* (>47%) were significantly more enriched in RNA microbial communities (p_Kruskal-Wallis_ < 0.05).

Genus-level analysis was performed on the *Actinobacteria*, *Firmicutes*, and *Proteobacteria* and among the 569 genera detected, 330, 323, and 309 genera showed significant differences between DNA and RNA microbial communities at N fertilization amount of 0, 224, and 448 kg N ha^-1^, respectively (Figure S4). Sequences associated with *Bacillus*, *Clostridium*, *Paenibacillus*, *Sporosarcina* of *Firmicutes* and *Bradyrhizobium*, *Methyloversatilis*, and *Nitrosomonas* of *Proteobacteria* more enriched in RNA than DNA. On the other hand, *Gaiella* from *Actinobacteria* and *Sphingomonas* of *Proteobacteria* were more abundant in DNA.

### Taxa associated with nitrogen cycle-related bacteria showed a short-term response since fertilization only in RNA microbial communities

In comparing pre- and post-fertilization soil samples, differences in soil microbial communities were observed only in RNA libraries (Table 1, Table S4). Taxa that were significantly different before and since fertilization were associated with 10 phyla (Figure 7). Additionally, these differences were only observed 69 days since fertilization, where the relative abundances of *Acidobacteria*, *Armatimonadetes*, *Firmicutes*, and *Planctomycetes* were increased compared to before fertilization, and the relative abundances of *Actinobacteria*, *Bacteroidetes*, *Chloroflexi*, and *Latescibacteria* decreased.

**Figure 7.**
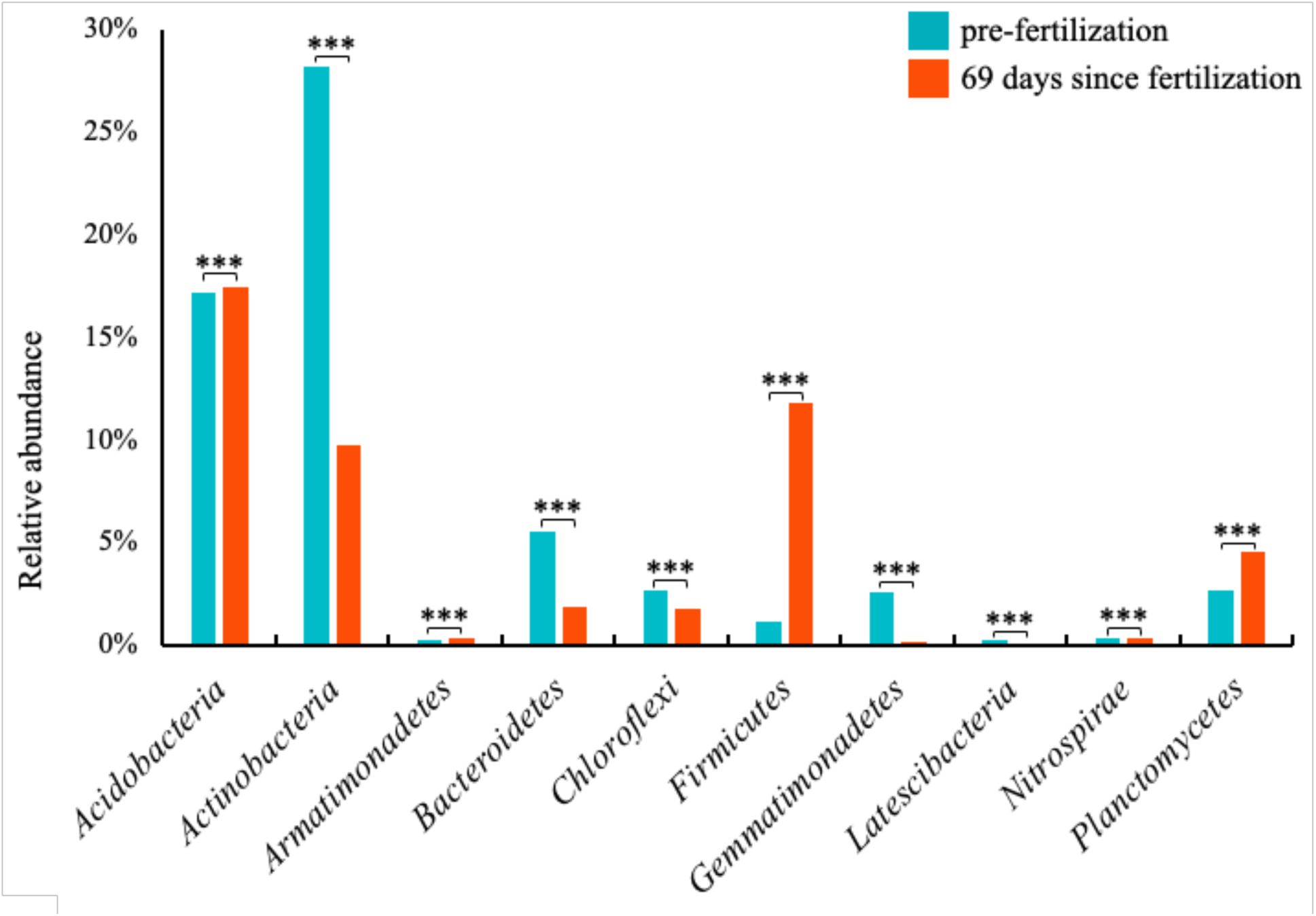
Phylum-level responses to time since fertilization in RNA microbial communities. The average relative abundances of phyla over time since fertilization were summarized. Letters “***” denote significant differences of relative abundance between pre- and 69 days since fertilization at a p-value < 0.05 as assess by Kruskal-Wallis with post hoc Dunn’s test.

The most enrichment since fertilization was observed in the *Firmicutes*, in which relative abundance was increased 10 folds, and *Planctomycetes* also increased by about 1.7 folds. These phyla are notable because they are known to contain known nitrogen cycling taxa (54, 55). To better explore the response to fertilization of nitrogen cycling taxa, we obtained taxa that are associated with nitrogen fixation, nitrification, and denitrification from the Fungene database. These taxa include 51 genera associated with *Firmicutes*, *Nitrospirae*, and *Planctomycetes*. We compared the differences of these genera between DNA and RNA libraries.

Overall, the total relative abundance of these genera comprised 1.18% and 8.51% in the DNA and RNA microbial communities, respectively (Figure 8). The large majority of these genera (with the exception of four genera) showed significant differences between DNA and RNA, and among them, *Bacillus, Paenibacillus*, and *Sporosarcina* were the most abundant (> 20%) in the RNA microbial community.

**Figure 8.**
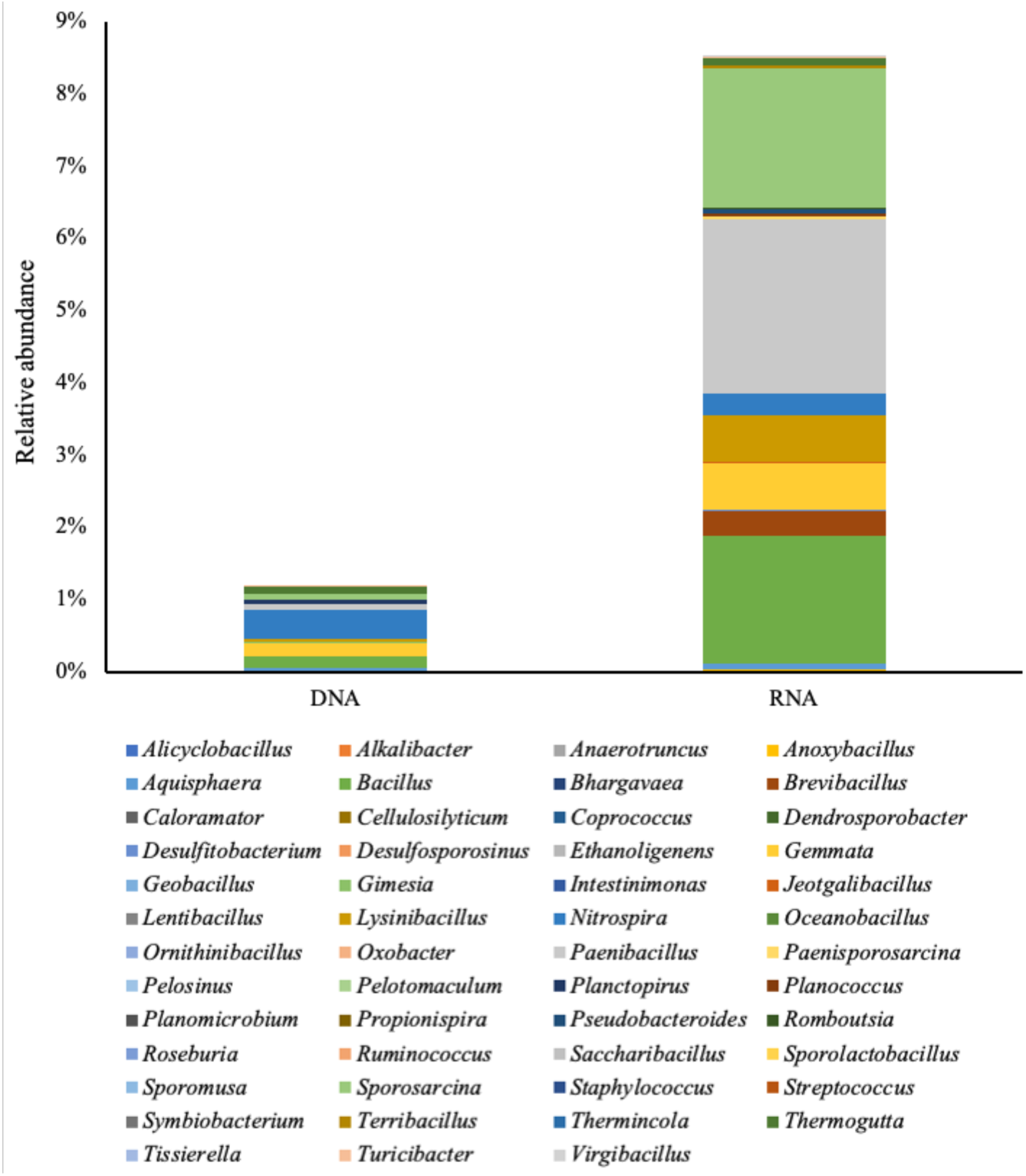
Comparison of nitrogen cycling-related bacteria in the DNA and RNA microbial communities. The average relative abundances of genus associated with nitrogen fixation, nitrification, and denitrification were summarized.

The taxa that showed distinct responses in RNA compared to DNA were classified by their known nitrogen cycling functions (excluding taxa with multiple functional annotations). Only taxa associated with denitrification in the RNA microbial communities showed a significant difference (Figure 9, Table S6) between pre- and post-fertilization, consistent with the observation that denitrifying bacteria were consistently enriched since fertilization (56).

**Figure 9.**
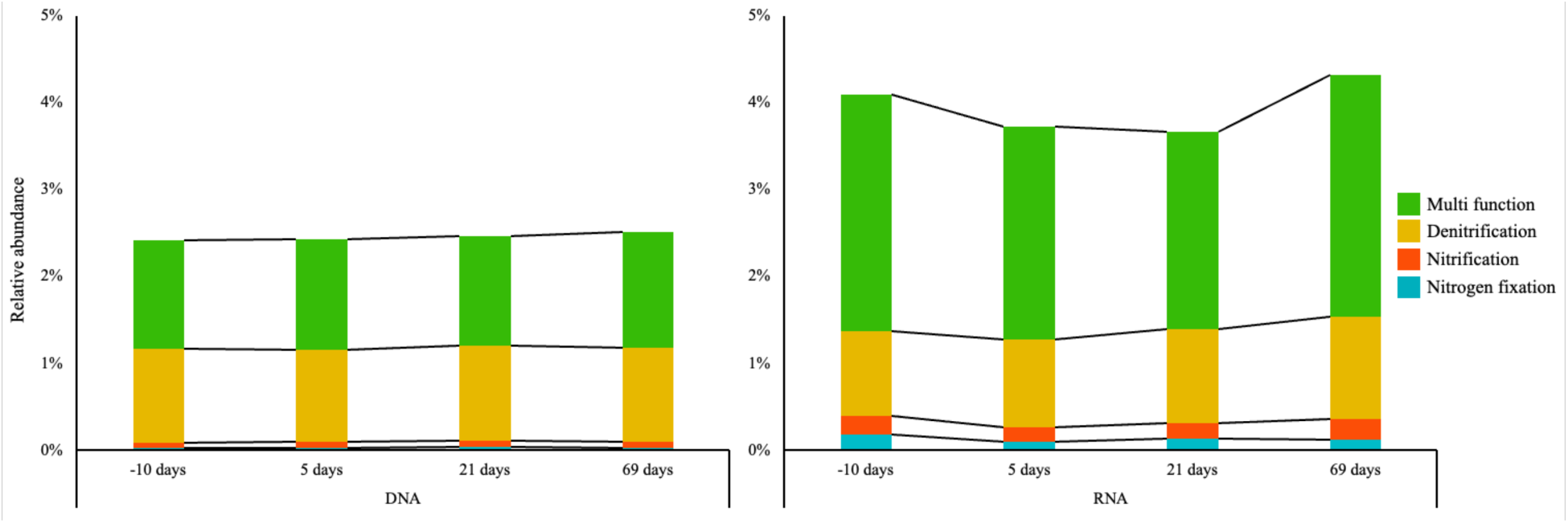
Comparison of nitrogen cycling-related bacteria in the DNA and RNA microbial communities according to time since fertilization. The average relative abundances of bacteria associated with nitrogen fixation, nitrification, and denitrification function were summarized.

## Discussion

In direct comparisons of *M.* x *giganteus* soil microbiomes from DNA and RNA extractions, we found that the most significant factor in explaining variation between microbiomes was its sequencing library of origin, even more so than experimental factors of stand age, N fertilization amount, or sampling day (Table S1). DNA and RNA microbiomes also had significantly different alpha diversity, with increased diversity and less variation observed in DNA relative to RNA. These results are consistent with what is known about DNA and RNA. DNA represents the potential genes or membership that may be active and thus is expected to represent more diverse membership with the potential to become metabolically active. RNA, which is actively transcribed, represents growing members, and its higher variability is consistent with its dynamic responses. Previous studies have shown that the RNA microbial community may also have lower alpha diversity because it does not contain the sequences of dormant or dead cells and also has greater variability in response to the environment (47, 57–59).

Overall, the large majority of membership between DNA and RNA was shared (greater than 90%), suggesting that both methods identify the similar presence of taxa. The abundance of these shared taxa, however, could be significantly different between DNA and RNA, and most of the shared taxa were more enriched in RNA. Based on the assumption that taxa observed in both methods are the most reliable, it is likely that DNA-based methods are underestimating the relative abundance of taxa. Further, these differences between DNA and RNA methods contributed to differences in estimated alpha diversity and varying observations of the microbial community response to plant host stand age and fertilization.

In response to both stand age and N fertilization amount, significant differences were observed in both DNA and RNA communities. While the overall pattern and ranking of differences were similar, the magnitude of this change and taxonomic membership driving these differences varied between DNA and RNA approaches. The most significant difference we observed in *M.* x *giganteus* soil microbial communities between the two library methods was in response to nitrogen fertilization. Only RNA microbial communities showed differences pre- and post-fertilization and only at day 69. RNA is able to show more rapid changes in response to changes in environmental conditions than DNA (60, 61), and here, we show the ability of RNA to capture a relatively short-term response over the course of one growing season in *M.* x *giganteus*, which is not observed in DNA. These results are consistent with RNA’s short half-life of several minutes to several hours (62) and also justify its usage for measuring short-term seasonal responses in bioenergy soils. Our results showed that it was not until over two months that a response different to pre-fertilization conditions was observed in RNA, providing some insight into the metabolic response of soil microbes to fertilization in these soils.

Among the taxa which were found to be uniquely identified in RNA libraries were members associated with nitrogen-cycling, including members of *Firmicutes*, *Nitrospirae*, and *Planctomycetes* which were enriched with both the presence of fertilizer and increasing nitrogen fertilizer. These results are consistent with previous studies which have shown that *Firmicutes* are enriched when nitrogen fertilizers are used (63–67). In the context of taxa associated with nitrogen cycling, it was confirmed that the RNA-based approach could better detect the denitrification function among the nitrogen cycling functions. This result is consistent with the results of previous studies that the application of nitrogen fertilizers suppressed the activity of nitrogen-fixing bacteria and enhanced denitrifying bacteria (56, 68). These results also emphasize that DNA may underestimate or miss the contribution of nitrogen-cycling taxa, which are highly relevant for nitrogen management in bioenergy systems. In addition to these taxa, we also found that members of dominant soil phyla, *Actinobacteria* and *Proteobacteria* are underestimated using DNA methods alone.

In summary, we found that DNA and RNA methods for characterizing the general response of microbial communities varied. With relevance to developing sustainable bioenergy crops and understanding the role of microbes in nutrient cycling, RNA appears to capture better the response of taxa known to be involved in nitrogen cycling and is also more sensitive to seasonal shifts in microbiomes. To better link microbial communities to ecosystem processes, we need to move towards characterizing the functional response of microbial communities. Due to costs, the first step in this characterization is often phylogenetic characterization of SSU genes based on DNA. Our results indicate that this method alone may bias against the composition results of the relevant microbial membership.

Notably, the integration of RNA-based methods into an experiment adds significant costs, requiring materials to quickly preserve samples for RNA extraction and typically more time for extraction and library preparation. RNA used for SSU characterization can be a complement to DNA-based studies, as it leverages the advantages and throughput of indicator gene amplification while not being as expensive as metatranscriptomics strategies. Based on our results, we recommended that DNA can be used for the initial and broad characterization of community membership. The use of RNA for SSU characterization could be used to complement DNA characterization when experimental questions have been developed. In the context of our experiment, DNA-based analyses were used to validate that there was a significant response to stand age and fertilization. RNA-based analyses were more helpful in identifying the specific taxa that respond to fertilization. With these specific taxa now identified, future research will be focused on functional characterization, guided by the result of this study (e.g., microbial responses to fertilization responses are most significant two months since fertilization). More broadly, in our understanding of microbial ecology, increasing numbers of studies are identifying the environments or gradients for which microbial communities are changing. In future work, it will be necessary to emphasize which taxa or what functions are changing, and our results indicate that RNA-based SSU characterization may be a substantial resource.

## Acknowledgments

This work was funded by the DOE Center for Advanced Bioenergy and Bioproducts Innovation (U.S. Department of Energy, Office of Science, Office of Biological and Environmental Research under Award Number DE-SC0018420). Any opinions, findings, and conclusions or recommendations expressed in this publication are those of the author(s) and do not necessarily reflect the views of the U.S. Department of Energy.

---

**Figure S1.**
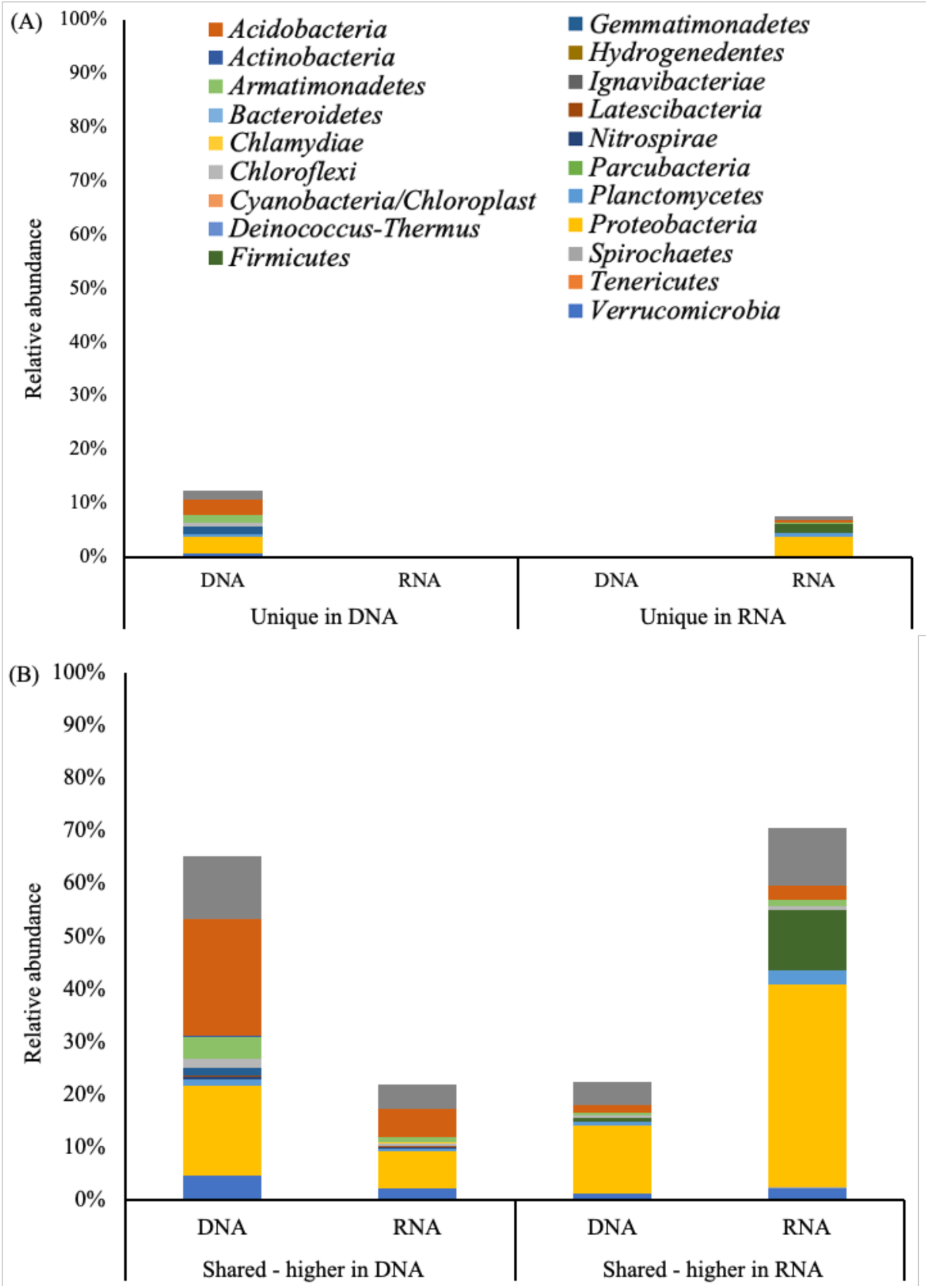
Phylum level differences in DNA and RNA microbial communities based on unique ASVs in DNA or RNA only (A) or shared ASVS and their enrichment in either DNA or RNA (B). Relative abundances of annotated ASVs are shown, identified to their closest match in the RDP classifier.

**Figure S2.**
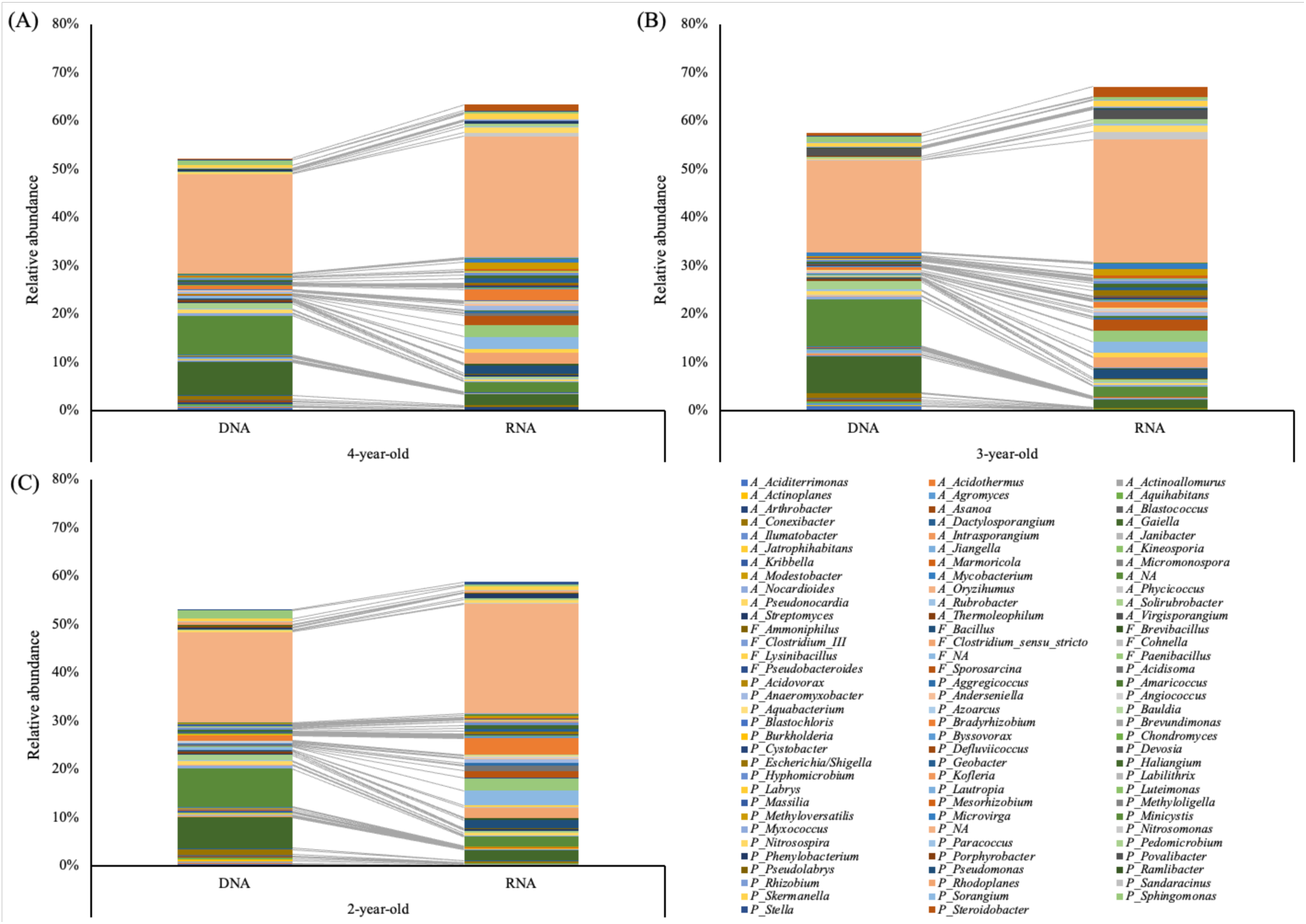
Comparison of dominant genus (> 0.1% relative abundance) in *Actinobacteria*, *Firmicutes*, and *Proteobacteria* between DNA and RNA microbial communities. (A) 4-year-old *M.* x *giganteus*, 73 genera. (B) 3-year-old *M.* x *giganteus*, 77 genera. (C) 2-year-old *M.* x *giganteus*, 74 genera. All genera included this analysis were significantly different between DNA and RNA microbial communities (p_Kruskal-Wallis_ < 0.05).

**Figure S3.**
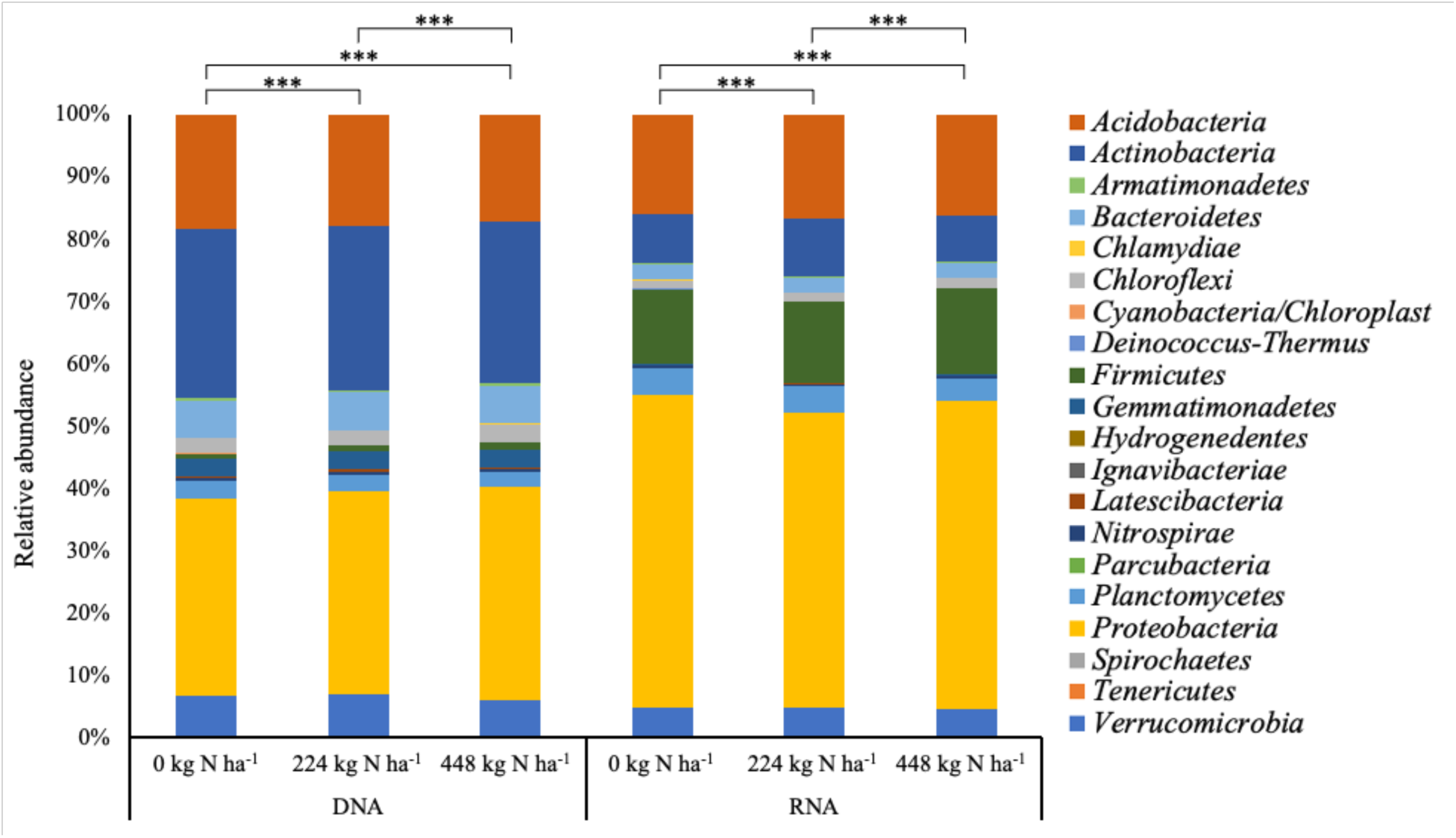
Phylum level differences in DNA and RNA microbial communities according to N fertilization amount differences. Relative abundances of annotated ASVs are shown, identified to their closest match in the RDP classifier. The letters “***” denote significant differences of relative abundance between different stand age of *M.* x *giganteus* at a p-value < 0.05 as assess by Kruskal-Wallis with post hoc Dunn’s test.

**Figure S4.**
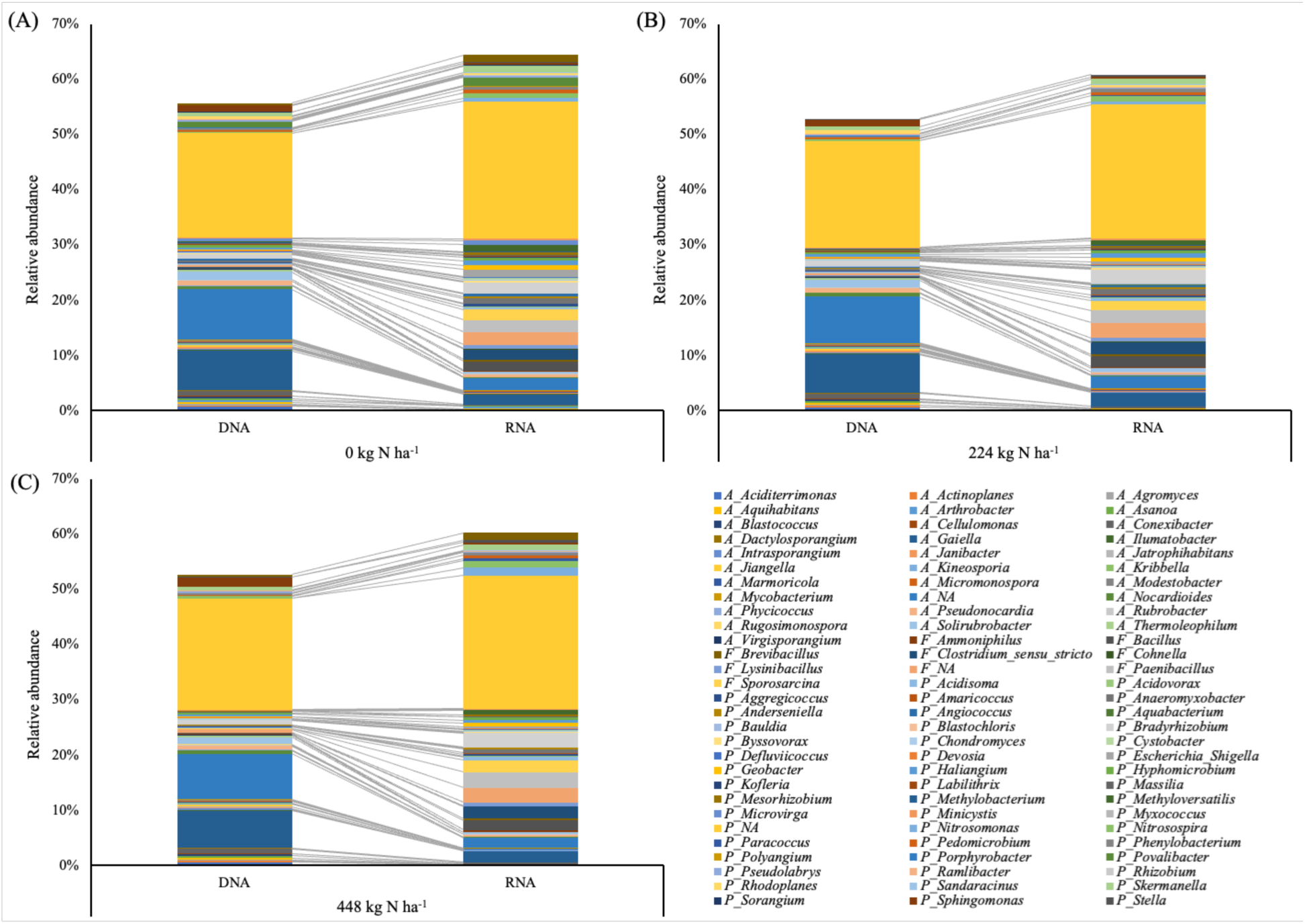
The dynamics of the 88 major genus (> 0.1% relative abundance) in the *Actinobacteria*, *Firmicutes*, and *Proteobacteria* between DNA and RNA microbial communities. (A) 0 kg N ha^-1^ of fertilizer applied *M.* x *giganteus* soil including 79 genera. (B) 224 kg N ha^-1^ of fertilizer applied *M.* x *giganteus* soil including 76 genera. (C) 448 kg N ha^-1^ of fertilizer applied *M.* x *giganteus* soil including 74 genera. All genera included this analysis were significantly different between DNA and RNA microbial communities (p_Kruskal-Wallis_ < 0.05).

**Table S1.** Comparison of microbial community dissimilarity using permutational multivariate analysis of variance (PERMANOVA) for bacterial communities in *M.* x *giganteus* soil samples.

| Response variable | Dissimilarity |
| --- | --- |
| Nucleic Acid<br>(DNA and RNA) | $R^2$ PERMANOVA: 0.117<br>$p$PERMANOVA: 0.001 |
| Stand ages<br>(2, 3, or 4-year-old) | $R^2$ PERMANOVA: 0.030<br>$p$PERMANOVA: 0.001 |
| N fertilization amount<br>(0, 224, and 448 kg N ha<sup>-1</sup>) | $R^2$ PERMANOVA: 0.008<br>$p$PERMANOVA: 0.001 |
| Time since fertilization<br>(-10, 5, 21, and 69 days) | $R^2$ PERMANOVA: 0.003<br>$p$PERMANOVA: 0.015 |
| Fertilization history<br>(fertilized or unfertilized) | $R^2$ PERMANOVA: 0.002<br>$p$PERMANOVA: n.s |

**Table S2.** Kruskal-Wallis with post hoc Dunn’s test comparing the average relative abundances of phyla between DNA and RNA microbial communities.

| Phylum | Average relative abundance |  | p-value |
| --- | --- | --- | --- |
|  | DNA | RNA |  |
| *Acidobacteria* | 17.88% | 16.29% | 6.54E-09 |
| *Actinobacteria* | 26.46% | 8.18% | 2.22E-87 |
| *Armatimonadetes* | 0.31% | 0.23% | 1.92E-17 |
| *Bacteroidetes* | 6.02% | 2.34% | 1.13E-77 |
| *Chlamydiae* | 0.05% | 0.10% | 1.60E-93 |
| *Chloroflexi* | 2.58% | 1.47% | 3.04E-90 |
| *Cyanobacteria/Chloroplast* | 0.02% | 0.03% | 7.67E-93 |
| *Deinococcus-Thermus* | 0.00% | 0.02% | 4.75E-07 |
| *Firmicutes* | 1.05% | 12.90% | 1.13E-92 |
| *Gemmatimonadetes* | 2.81% | 0.14% | 1.33E-87 |
| *Hydrogenedentes* | 0.00% | 0.00% | 7.75E-83 |
| *Ignavibacteriae* | 0.01% | 0.01% | 2.61E-11 |
| *Latescibacteria* | 0.28% | 0.10% | 4.30E-83 |
| *Nitrospirae* | 0.45% | 0.33% | 1.69E-74 |
| *Parcubacteria* | 0.05% | 0.00% | 7.05E-68 |
| *Planctomycetes* | 2.64% | 4.06% | 4.49E-98 |
| *Proteobacteria* | 32.76% | 48.97% | 3.08E-90 |
| *Spirochaetes* | 0.01% | 0.01% | 1.50E-94 |
| *Tenericutes* | 0.00% | 0.00% | 1.93E-33 |
| *Verrucomicrobia* | 6.63% | 4.81% | 3.18E-15 |

**Table S3.** Pairwise permutational multivariate analysis of variance (PERMANOVA) for comparing the effect of stand age and fertilization on the DNA and RNA microbial community dissimilarity.

| Response variable | Stand age |  | N fertilization amount (kg N ha<sup>-1</sup>) |  |
| --- | --- | --- | --- | --- |
| DNA microbial community | 4-year-old vs 3-year-old | $R^2_{\text{pairwisePERMANOVA}} = 0.063$<br>$p_{\text{pairwisePERMANOVA}} = 0.001$ | 0 vs 224 | $R^2_{\text{pairwisePERMANOVA}} = 0.014$<br>$p_{\text{pairwisePERMANOVA}} = 0.023$ |
| | 4-year-old vs 2-year-old | $R^2_{\text{pairwisePERMANOVA}} = 0.073$<br>$p_{\text{pairwisePERMANOVA}} = 0.001$ | 0 vs 448 | $R^2_{\text{pairwisePERMANOVA}} = 0.021$<br>$p_{\text{pairwisePERMANOVA}} = 0.018$ |
| | 3-year-old vs 2-year-old | $R^2_{\text{pairwisePERMANOVA}} = 0.210$<br>$p_{\text{pairwisePERMANOVA}} = 0.001$ | 224 vs 448 | $R^2_{\text{pairwisePERMANOVA}} = 0.015$<br>$p_{\text{pairwisePERMANOVA}} = 0.023$ |
| RNA microbial community | 4-year-old vs 3-year-old | $R^2_{\text{pairwisePERMANOVA}} = 0.039$<br>$p_{\text{pairwisePERMANOVA}} = 0.001$ | 0 vs 224 | $R^2_{\text{pairwisePERMANOVA}} = 0.011$<br>$p_{\text{pairwisePERMANOVA}} = 0.031$ |
| | 4-year-old vs 2-year-old | $R^2_{\text{pairwisePERMANOVA}} = 0.057$<br>$p_{\text{pairwisePERMANOVA}} = 0.001$ | 0 vs 448 | $R^2_{\text{pairwisePERMANOVA}} = 0.013$<br>$p_{\text{pairwisePERMANOVA}} = 0.029$ |
| | 3-year-old vs 2-year-old | $R^2_{\text{pairwisePERMANOVA}} = 0.152$<br>$p_{\text{pairwisePERMANOVA}} = 0.001$ | 224 vs 448 | $R^2_{\text{pairwisePERMANOVA}} = 0.012$<br>$p_{\text{pairwisePERMANOVA}} = 0.029$ |

**Table S4.**
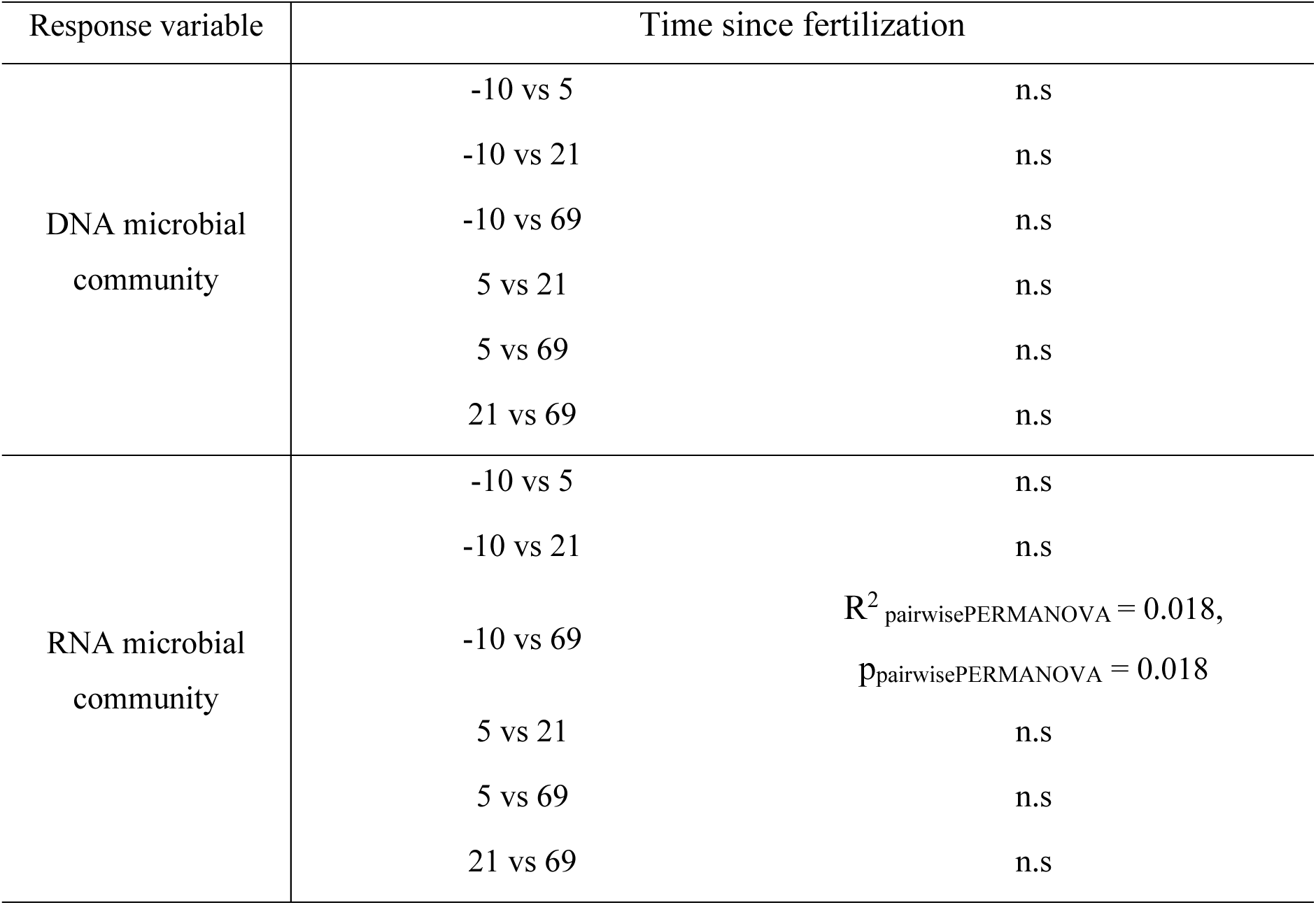
Pairwise permutational multivariate analysis of variance (PERMANOVA) for comparing the effect of time since fertilization on the DNA and RNA microbial community dissimilarity.

| Response variable | Time since fertilization |  |
| --- | --- | --- |
| DNA microbial community | -10 vs 5 | n.s |
|  | -10 vs 21 | n.s |
|  | -10 vs 69 | n.s |
|  | 5 vs 21 | n.s |
|  | 5 vs 69 | n.s |
|  | 21 vs 69 | n.s |
| RNA microbial community | -10 vs 5 | n.s |
|  | -10 vs 21 | n.s |
| | -10 vs 69 | $R^2_{\text{pairwisePERMANOVA}} = 0.018,$<br>$p_{\text{pairwisePERMANOVA}} = 0.018$ |
|  | 5 vs 21 | n.s |
|  | 5 vs 69 | n.s |
|  | 21 vs 69 | n.s |

**Table S5.** Kruskal-Wallis with post hoc Dunn’s test comparing the average relative abundances of phyla between DNA and RNA microbial communities by different N fertilization. Denote N0, N224, and N448 are N fertilization of 0 kg N ha^-1^, 224 kg N ha^-1^, and 448 kg N ha^-1^, respectively.

|  | Phylum | Difference in average relative abundance |  | p-value |
| --- | --- | --- | --- | --- |
| DNA microbial community | *Acidobacteria* | 18.44% | 17.23% | 3.62E-02<br>between N0 and N448 |
|  | *Chloroflexi* | 2.30% | 2.90% | 1.24E-02<br>between N224 and N448 |
|  | *Latescibacteria* | 0.32% | 0.30% | 1.79E-04<br>between N0 and N448 |
|  |  | 0.30% | 0.21% | 1.30E-03<br>between N224 and N448 |
|  | *Proteobacteria* | 31.54% | 34.10% | 3.17E-04<br>between N0 and N448 |
| RNA microbial community | *Actinobacteria* | 9.18% | 7.39% | 1.33E-03<br>between N224 and N448 |
|  | *Firmicutes* | 12.91% | 13.78% | 2.55E-03<br>between N224 and N448 |
|  | *Gemmatimonadetes* | 0.12% | 0.13% | 4.52E-02<br>between N0 and N448 |
|  | *Hydrogenedentes* | 0.0054% | 0.0047% | 2.47E-02<br>between N0 and N224 |
|  |  | 0.0047% | 0.0035% | 2.62E-02<br>between N224 and N448 |
|  | *Latescibacteria* | 0.10% | 0.08% | 3.87E-03<br>between N224 and N448 |
|  | *Nitrospirae* | 0.30% | 0.31% | 1.59E-04<br>between N0 and N448 |
|  | *Proteobacteria* | 50.08% | 49.46% | 3.27E-05<br>between N0 and N448 |

**Table S6.**
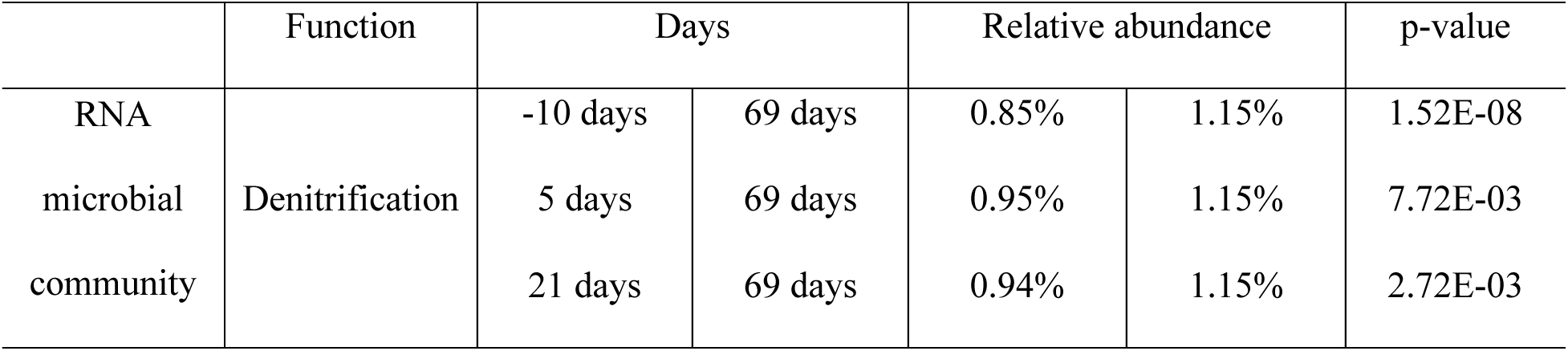
Kruskal-Wallis with post hoc Dunn’s test comparing the average relative abundances of nitrogen cycling functions in RNA microbial communities by time since fertilization.

## Reference

1. Lewandowski I, Clifton-Brown JC, Scurlock JMO, Huisman W. 2000. Miscanthus: European experience with a novel energy crop. Biomass and Bioenergy 19:209–227.

2. Heaton EA, Dohleman FG, Miguez AF, Juvik JA, Lozovaya V, Widholm J, Zabotina OA, McIsaac GF, David MB, Voigt TB, Boersma NN, Long SP. 2010. Chapter 3 - Miscanthus: A Promising Biomass Crop, p. 75–137. In Kader, J-C, Delseny, M (eds.), . Academic Press.

3. Jørgensen U. 2011. Benefits versus risks of growing biofuel crops: The case of Miscanthus. Current Opinion in Environmental Sustainability.

4. Brosse N, Dufour A, Meng X, Sun Q, Ragauskas A. 2012. Miscanthus: A fast-growing crop for biofuels and chemicals production. Biofuels, Bioproducts and Biorefining 6:580–598.

5. Heaton EA, Dohleman FG, Long SP. 2008. Meeting US biofuel goals with less land: The potential of Miscanthus. Global Change Biology 14:2000–2014.

6. Boehmel C, Lewandowski I, Claupein W. 2008. Comparing annual and perennial energy cropping systems with different management intensities. Agricultural Systems 96:224–236.

7. Lewandowski I, Schmidt U. 2006. Nitrogen, energy and land use efficiencies of miscanthus, reed canary grass and triticale as determined by the boundary line approach. Agriculture, Ecosystems and Environment 112:335–346.

8. Schwarz H, Liebhard P, Ehrendorfer K, Ruckenbauer P. 1994. The effect of fertilization on yield and quality of Miscanthus sinensis “Giganteus.” Industrial Crops and Products 2:153–159.

9. Lewandowski I, Heinz A. 2003. Delayed harvest of miscanthus - Influences on biomass quantity and quality and environmental impacts of energy production. European Journal of Agronomy 19:45–63.

10. Di HJ, Cameron KC. 2002. Nitrate leaching in temperate agroecosystems: Sources, factors and mitigating strategies. Nutrient Cycling in Agroecosystems 64:237–256.

11. Christian DG, Riche AB. 1998. Nitrate leaching losses under Miscanthus grass planted on a silty clay loam soil. Soil Use and Management 14:131–135.

12. Heaton EA, Clifton-Brown J, Voigt TB, Jones MB, Long SP. 2004. Miscanthus for renewable energy generation: European Union experience and projections for Illinois. Mitigation and Adaptation Strategies for Global Change 9:433–451.

13. Styles D, Thorne F, Jones MB. 2008. Energy crops in Ireland: An economic comparison of willow and Miscanthus production with conventional farming systems. Biomass and Bioenergy 32:407–421.

14. Witzel CP, Finger R. 2016. Economic evaluation of Miscanthus production - A review. Renewable and Sustainable Energy Reviews 53:681–696.

15. Arnoult S, Brancourt-Hulmel M. 2015. A Review on Miscanthus Biomass Production and Composition for Bioenergy Use: Genotypic and Environmental Variability and Implications for Breeding. Bioenergy Research 8:502–526.

16. Clifton-Brown JC, Lewandowski I, Andersson B, Basch G, Christian DG, Kjeldsen JB, Jørgensen U, Mortensen J v., Riche AB, Schwarz KU, Tayebi K, Teixeira F. 2001. Performance of 15 Miscanthus genotypes at five sites in Europe. Agronomy Journal 93:1013–1019.

17. McIsaac GF, David MB, Mitchell CA. 2010. Miscanthus and Switchgrass Production in Central Illinois: Impacts on Hydrology and Inorganic Nitrogen Leaching. Journal of Environmental Quality 39:1790–1799.

18. Cosentino SL, Patanè C, Sanzone E, Copani V, Foti S. 2007. Effects of soil water content and nitrogen supply on the productivity of Miscanthus × giganteus Greef et Deu. in a Mediterranean environment. Industrial Crops and Products 25:75–88.

19. Christian DG, Riche AB, Yates NE. 2008. Growth, yield and mineral content of Miscanthus × giganteus grown as a biofuel for 14 successive harvests. Industrial Crops and Products 28:320–327.

20. Jezowski S. 2008. Yield traits of six clones of Miscanthus in the first 3 years following planting in Poland. Industrial Crops and Products 27:65–68.

21. Jezowski S, Głowacka K, Kaczmarek Z. 2011. Variation on biomass yield and morphological traits of energy grasses from the genus Miscanthus during the first years of crop establishment. Biomass and Bioenergy 35:814–821.

22. Amougou N, Bertrand I, Machet JM, Recous S. 2011. Quality and decomposition in soil of rhizome, root and senescent leaf from Miscanthus x giganteus, as affected by harvest date and N fertilization. Plant and Soil 338:83–97.

23. Gauder M, Graeff-Hönninger S, Lewandowski I, Claupein W. 2012. Long-term yield and performance of 15 different Miscanthus genotypes in southwest Germany. Annals of Applied Biology 160:126–136.

24. McLaughlin SB, Kszos LA. 2005. Development of switchgrass (Panicum virgatum) as a bioenergy feedstock in the United States. Biomass and Bioenergy 28:515–535.

25. Lesur C, Jeuffroy MH, Makowski D, Riche AB, Shield I, Yates N, Fritz M, Formowitz B, Grunert M, Jorgensen U, Laerke PE, Loyce C. 2013. Modeling long-term yield trends of Miscanthus×giganteus using experimental data from across Europe. Field Crops Research 149:252–260.

26. Thompson KA, Deen B, Dunfield KE. 2016. Soil denitrifier community size changes with land use change to perennial bioenergy cropping systems. SOIL 2:523–535.

27. Li D, Voigt TB, Kent AD. 2016. Plant and soil effects on bacterial communities associated with Miscanthus × giganteus rhizosphere and rhizomes. GCB Bioenergy 8:183–193.

28. Tejera M, Boersma N, Vanloocke A, Archontoulis S, Dixon P, Miguez F, Heaton E. 2019. Multi-year and Multi-site Establishment of the Perennial Biomass Crop Miscanthus × giganteus Using a Staggered Start Design to Elucidate N Response. Bioenergy Research 12:471–483.

29. Miguez FE, Villamil MB, Long SP, Bollero GA. 2008. Meta-analysis of the effects of management factors on Miscanthus × giganteus growth and biomass production. Agricultural and Forest Meteorology 148:1280–1292.

30. Cadoux S, Riche AB, Yates NE, Machet JM. 2012. Nutrient requirements of Miscanthus x giganteus: Conclusions from a review of published studies. Biomass and Bioenergy 38:14–22.

31. Wang D, Maughan MW, Sun J, Feng X, Miguez F, Lee D, Dietze MC. 2012. Impact of nitrogen allocation on growth and photosynthesis of Miscanthus (Miscanthus × giganteus). GCB Bioenergy 4:688–697.

32. Arundale RA, Dohleman FG, Voigt TB, Long SP. 2014. Nitrogen Fertilization Does Significantly Increase Yields of Stands of Miscanthus × giganteus and Panicum virgatum in Multiyear Trials in Illinois. Bioenergy Research 7:408–416.

33. Lee MS, Wycislo A, Guo J, Lee DK, Voigt T. 2017. Nitrogen fertilization effects on biomass production and yield components of miscanthus × giganteus. Frontiers in Plant Science 8:544.

34. Finnan J, Burke B. 2016. Nitrogen fertilization of Miscanthus x giganteus: effects on nitrogen uptake, growth, yield and emissions from biomass combustion. Nutrient Cycling in Agroecosystems 106:249–256.

35. Monti A, Zegada-Lizarazu W, Zanetti F, Casler M. 2019. Nitrogen Fertilization Management of Switchgrass, Miscanthus and Giant Reed: A Review, p. 87–119. In Advances in Agronomy.

36. Keymer DP, Kent AD. 2014. Contribution of nitrogen fixation to first year Miscanthus × giganteus. GCB Bioenergy 6:577–586.

37. Davis SC, Parton WJ, Dohleman FG, Smith CM, del Grosso S, Kent AD, DeLucia EH. 2010. Comparative biogeochemical cycles of bioenergy crops reveal nitrogen-fixation and low greenhouse gas emissions in a Miscanthus × giganteus agro-ecosystem. Ecosystems 13:144–156.

38. Mao Y, Yannarell AC, Davis SC, Mackie RI. 2013. Impact of different bioenergy crops on N-cycling bacterial and archaeal communities in soil. Environmental Microbiology 15:928–942.

39. Jiaron Guo. 2016. Rhizosphere metagenomics of three biofuel crops.

40. Liu Y, Ludewig U. 2019. Nitrogen-dependent bacterial community shifts in root, rhizome and rhizosphere of nutrient-efficient Miscanthus x giganteus from long-term field trials. GCB Bioenergy 11:1334–1347.

41. Ma L, Rocha FI, Lee J, Choi J, Tejera M, Sooksa-Nguan T, Boersma N, VanLoocke A, Heaton E, Howe A. 2021. The impact of stand age and fertilization on the soil microbiome of miscanthus × giganteus. Phytobiomes Journal 5:51–59.

42. Papp K, Hungate BA, Schwartz E. 2018. Microbial rRNA synthesis and growth compared through quantitative stable isotope probing with H218O. Applied and Environmental Microbiology 84:e02441–17.

43. Dlott G, Maul JE, Buyer J, Yarwood S. 2015. Microbial rRNA: RDNA gene ratios may be unexpectedly low due to extracellular DNA preservation in soils. Journal of Microbiological Methods 115:112–120.

44. Broughton LC, Gross KL. 2000. Patterns of diversity in plant and soil microbial communities along a productivity gradient in a Michigan old-field. Oecologia 125:420–427.

45. Freedman ZB, Romanowicz KJ, Upchurch RA, Zak DR. 2015. Differential responses of total and active soil microbial communities to long-term experimental N deposition. Soil Biology and Biochemistry 90:275–282.

46. Lennon JT, Jones SE. 2011. Microbial seed banks: the ecological and evolutionary implications of dormancy. Nature Reviews Microbiology 9:119–130.

47. Baldrian P, Kolařík M, Štursová M, Kopecký J, Valášková V, Větrovský T, Žifčáková L, Šnajdr J, Rídl J, Vlček Č, Voříšková J. 2012. Active and total microbial communities in forest soil are largely different and highly stratified during decomposition. The ISME Journal 6:248–258.

48. Zhang Y, Zhao Z, Dai M, Jiao N, Herndl GJ. 2014. Drivers shaping the diversity and biogeography of total and active bacterial communities in the South China Sea. Molecular Ecology 23:2260–2274.

49. Videira SS, Pereira e Silva M de C, de Souza Galisa P, Dias ACF, Nissinen R, Divan VLB, van Elsas JD, Baldani JI, Salles JF. 2013. Culture-independent molecular approaches reveal a mostly unknown high diversity of active nitrogen-fixing bacteria associated with Pennisetum purpureum—a bioenergy crop. Plant and Soil 373:737–754.

50. Studt JE, McDaniel MD, Tejera MD, VanLoocke A, Howe A, Heaton EA. 2021. Soil net nitrogen mineralization and leaching under Miscanthus × giganteus and Zea mays. GCB Bioenergy 13:1545–1560.

51. Chowdary D, Lathrop J, Skelton J, Curtin K, Briggs T, Zhang Y, Yu J, Wang Y, Mazumder A. 2006. Prognostic gene expression signatures can be measured in tissues collected in RNAlater preservative. Journal of Molecular Diagnostics 8:31–39.

52. Parada AE, Needham DM, Fuhrman JA. 2016. Every base matters: Assessing small subunit rRNA primers for marine microbiomes with mock communities, time series and global field samples. Environmental Microbiology 18:1403–14.

53. Apprill A, Mcnally S, Parsons R, Weber L. 2015. Minor revision to V4 region SSU rRNA 806R gene primer greatly increases detection of SAR11 bacterioplankton. Aquatic Microbial Ecology 75:129–137.

54. Delmont TO, Quince C, Shaiber A, Esen ÖC, Lee STM, Rappé MS, McLellan SL, Lücker S, Eren AM. 2018. Nitrogen-fixing populations of Planctomycetes and Proteobacteria are abundant in surface ocean metagenomes. Nature Microbiology 3:804–813.

55. Ren C, Zhang W, Zhong ZK, Han X, Yang G, Feng Y, Ren G. 2018. Differential responses of soil microbial biomass, diversity, and compositions to altitudinal gradients depend on plant and soil characteristics. Science of the Total Environment 610–611:750–758.

56. Knowles R. 1982. Denitrification. Microbiological reviews 46:43–70.

57. Li R, Tun HM, Jahan M, Zhang Z, Kumar A, Fernando D, Farenhorst A, Khafipour E. 2017. Comparison of DNA-, PMA-, and RNA-based 16S rRNA Illumina sequencing for detection of live bacteria in water. Scientific Reports 7:5752.

58. de Vrieze J, Pinto AJ, Sloan WT, Ijaz UZ. 2018. The active microbial community more accurately reflects the anaerobic digestion process: 16S rRNA (gene) sequencing as a predictive tool. Microbiome 6:63.

59. Prussin AJ, Torres PJ, Shimashita J, Head SR, Bibby KJ, Kelley ST, Marr LC. 2019. Seasonal dynamics of DNA and RNA viral bioaerosol communities in a daycare center. Microbiome 7:53.

60. Bulow FJ. 1970. RNA–DNA Ratios as Indicators of Recent Growth Rates of a Fish. Journal of the Fisheries Research Board of Canada 27:2343–2349.

61. Eichler S, Christen R, Höltje C, Westphal P, Bötel J, Brettar I, Mehling A, Höfle MG. 2006. Composition and dynamics of bacterial communities of a drinking water supply system as assessed by RNA- and DNA-based 16S rRNA gene fingerprinting. Applied and Environmental Microbiology 72:1858–1872.

62. Deana A, Belasco JG. 2005. Lost in translation: The influence of ribosomes on bacterial mRNA decay. Genes and Development 19:2526–2533.

63. Smercina DN, Evans SE, Friesen ML, Tiemann LK. 2019. To fix or not to fix: Controls on free-living nitrogen fixation in the rhizosphere. Applied and Environmental Microbiology 85:e02546–18.

64. Li JG, Shen MC, Hou JF, Li L, Wu JX, Dong YH. 2016. Effect of different levels of nitrogen on rhizosphere bacterial community structure in intensive monoculture of greenhouse lettuce. Scientific Reports 6:25305.

65. Zhou J, Guan D, Zhou B, Zhao B, Ma M, Qin J, Jiang X, Chen S, Cao F, Shen D, Li J. 2015. Influence of 34-years of fertilization on bacterial communities in an intensively cultivated black soil in northeast China. Soil Biology and Biochemistry 90:42–51.

66. Wagner M, Horn M. 2006. The Planctomycetes, Verrucomicrobia, Chlamydiae and sister phyla comprise a superphylum with biotechnological and medical relevance. Current Opinion in Biotechnology 17:241–249.

67. Leff JW, Jones SE, Prober SM, Barberán A, Borer ET, Firn JL, Harpole WS, Hobbie SE, Hofmockel KS, Knops JMH, McCulley RL, la Pierre K, Risch AC, Seabloom EW, Schütz M, Steenbock C, Stevens CJ, Fierer N. 2015. Consistent responses of soil microbial communities to elevated nutrient inputs in grasslands across the globe. Proceedings of the National Academy of Sciences of the United States of America 112:10967–10972.

68. Thompson KA, Deen B, Dunfield KE. 2018. Impacts of surface-applied residues on N-cycling soil microbial communities in miscanthus and switchgrass cropping systems. Applied Soil Ecology 130:79–83.

